# Bioengineering of a Chimeric *Metarhizium anisopliae* cPr1A Protease with Enhanced Binding and Enzyme Activity Through C-terminal Fusion of *Bombyx mori* Chitin-Binding Domain

**DOI:** 10.64898/2026.07.24.740453

**Authors:** Neha Maurya, Gurvinder Kaur Saini

**Affiliations:** Indian Institute of Technology Guwahati, Assam, India

## Abstract

*Metarhizium anisopliae* is an important entomopathogenic fungi used in biological control of agricultural pests, but its commercial application is limited by relatively slow host mortality. This study aimed to engineer a chimeric protease (cPr1A) with enhanced binding affinity and protease activity against insect cuticle. We hypothesized that stronger cuticle binding would increase local enzyme concentration at the cuticle surface and thereby enhance cuticle degradation. To achieve this, the *Bombyx mori* chitin-binding domain (BmCBD) was fused to the C-terminus of the Pr1A protease from *M. anisopliae*. Recombinant Pr1A and cPr1A were expressed in *Escherichia coli*, purified by Ni-NTA affinity chromatography. Binding and protease activity were assayed in triplicate using Samia ricini cuticle powder as substrate. Results are presented as mean +/- SEM. The chimeric protease cPr1A showed a 28.9% increase in cuticle binding compared to wild-type Pr1A (15.81 +/- 1.97 vs. 12.27 +/- 2.13 μg bound protein/mg cuticle powder; p < 0.002) and a 35% increase in protease activity (0.343 +/- 0.08 U/mg vs. 0.254 +/- 0.06 U/mg; p < 0.03). These results indicate that cPr1A is a promising candidate for overexpression in *M. anisopliae* to enhance cuticle degradation and potentially improve fungal virulence against insect pests.

## 1. Introduction

Chemical pesticides pose significant hazards to the environment. Eco-friendly fungal biopesticides have emerged as viable substitutes [1–3]. Entomopathogenic (EMP) fungus, *Metarhizium anisopliae*, offers significant promise as biological control agent for agricultural pests and human disease vectors [4–5]. *Metarhizium anisopliae* is an ascomycete that thrives globally. In agricultural environments, the diverse survival strategy of *Metarhizium* includes its persistence in soil as a saprophyte, colonization of the rhizosphere, endophytic growth, and pathogenic infection of insects, eventually producing green cylindrical conidia during sporulation on insect cadaver [6–9].

*M. anisopliae* is a well-studied fungus that has been approved as safe by the US EPA [10]. Unlike bacterial or viral biopesticides, it infects without ingestion and has been used successfully to control pests such as mosquitoes, ticks, and termites [11–14], but the slow killing speed of these mycoinsecticides, which can take days to weeks, limits their application and renders them less competitive than fast-acting chemical insecticides [15]. It utilizes mechanical pressure developed by the appressorium as well as a variety of hydrolases, including chitinase (Chi2), subtilisin-like proteases (Pr1), trypsin-like proteases (Pr2), and lipases to breach the insect cuticle [16]. Pr1A protease is a hallmark of the infection process and digests the protein components of the insect host’s cuticle, facilitating cuticle penetration, host colonization, destruction of insect tissues, and utilization of cuticular components as nutrient sources [17–18].

Genomic studies of *M. anisopliae* have revealed 11 subtilisin-like genes, Pr1A to Pr1K. Of these, Pr1A represents a key target for engineering biopesticides because of its high expression during cuticle penetration and its contribution to fungal virulence [16]. Enhanced binding affinity of the protease to the cuticle improves protease efficiency and accelerates cuticle degradation. For example, fusion of the *Bombyx mori* chitin-binding domain (BmChBD) to the subtilisin-like protease CDEP-1 of an insect-pathogenic fungi *Beauveria bassiana* significantly improved insecticidal activity against the aphid *Myzus persicae* larvae, resulting in increased virulence [19]. Pr1A, a serine protease, a key virulence-determining factor in the entomopathogenic fungus *Metarhizium anisopliae*, is crucial for the hydrolysis of the insect host cuticular protein matrix. The insect cuticle primarily consists of chitin fibrils embedded in the protein matrix. Increasing the binding affinity of Pr1A protease with the protein-chitin matrix of the insect host cuticle would improve protease-cuticle interaction by improving the targeting of the chimeric protease to the chitin component, which will bring the protease into closer with the cuticle, accelerate the proteolytic degradation of cuticular proteins, and increase fungal virulence. Therefore, we hypothesized that fusing the *Bombyx mori* chitin-binding domain (BmChBD) to the C-terminal of *M. anisopliae* Pr1A will result in a chimeric protease, cPr1A, with a higher binding affinity and increased cuticle-degrading potential.

## 2. Materials and methods

### 2.1 Bacterial Strains Reagents and Kits

#### 2.1(a) Bacterial Strains and plasmids

*E. coli* TOP10 cells were used for the cloning and maintenance of the pET- 28a(+) vector. *E. coli* BL21(DE3) cells (New England Biolabs, USA) were used for the expression of Pr1A and cPr1A proteins.

#### 2.1(b) PCR and Cloning

All cloning and expression experiments for Pr1A and cPr1A were done using the pET-28a(+) vector. PCR amplification of Pr1A and cPr1A genes was performed using Phusion DNA polymerase (Thermo Scientific), and PCR confirmation of positive clones was done using Taq DNA polymerase (New England Biolabs, USA). Cloning was done using the laboratory prepared Gibson assembly master mix **(Supplementary Table S1)**. NcoI (New England Biolabs, USA) and XhoI (New England Biolabs, USA) restriction enzymes were used for the linearization of the pET- 28a(+) vector. Primers for the amplification of Pr1A and BmChBD fragments were synthesized by GCC Biotech INDIA Pvt Ltd, India. The amplified genes and fragments were purified from the agarose gel using NucleoSpin Gel and PCR Clean-up kit (Macherey–Nagel, Duren, Germany).

#### 2.1(c) Chemicals & Reagents

Luria-Bertani (LB) broth, sodium hydroxide (NaOH), kanamycin, Tris base, Bradford reagent, disodium ethylenediaminetetraacetate (EDTA), and Coomassie Brilliant Blue, were obtained from HiMedia Laboratories (India). Sodium dodecyl sulfate (SDS) and 3,5-Dinitrosalicylic acid (DNSA) were acquired from SRL (India). DNA and protein ladders were purchased from Thermo Scientific and Bio-Rad, India, respectively.

### 2.2 Protease Induction and Production in Cultured *Metarhizium anisopliae*

Conidia of *M. anisopliae* (1×10^6^ conidia/mL) were inoculated into Sabouraud dextrose broth (SDB; 2% dextrose, 1% peptone, pH 5.6) and incubated in an orbital incubator shaker at 28°C, 180 rpm for 48 h to obtain actively growing mycelia. The mycelia were harvested by filtration through Whatman No.1 filter paper, washed thoroughly with sterile distilled water, and transferred to protease induction medium (PIM; KH_2_PO_4_ 0.1%, MgSO_4_·7H_2_O 0.05%, Colloidal chitin 1%) to induce expression of Pr1A protease. Mycelia were subsequently harvested from PIM by filtration, washed with sterile distilled water, flash-frozen in liquid nitrogen, and stored at -80°C until RNA isolation.

### 2.3 RNA Isolation, cDNA Synthesis, and Pr1A Gene Amplification

The total RNA was isolated from PIM-induced mycelia using the RNAiso Plus kit (Takara, Japan) according to the manufacturer’s protocol. Residual genomic DNA was removed by DNase I treatment (Sigma-Aldrich). RNA concentration was quantified using a NanoDrop spectrophotometer, and its integrity was verified by 1% agarose gel electrophoresis by visualizing the presence of intact 28S and 18S rRNA bands. One microgram of total RNA was converted into cDNA using the SuperScript™ III First-Strand Synthesis System (Invitrogen, USA). The synthesized cDNA was used as a template for the amplification of the Pr1A gene using gene-specific primers.

### 2.4 Amplification and Cloning of the Wild-Type Pr1A Gene

A set of gene-specific forward and reverse primers were designed based on the Pr1A gene sequence (NCBI Accession No. M73795.1). The amplification of Pr1A gene was done using primers and cDNA as the template. PCR cycling conditions and primer sequences and are provided in **(Supplementary Table S2 and S3)**. The Pr1A amplicon and NcoI/XhoI-digested pET-28a(+) plasmid were resolved on a 1% agarose gel and purified from the gel using the NucleoSpin Gel and PCR Clean-up kit. The purified Pr1A insert with complementary vector overhangs was cloned into the linearized pET-28a(+) vector using the lab-prepared Gibson assembly mix at 50°C for 60 min. The Gibson assembly mix (10 μL) following incubation was transformed into 50 μL of chemically competent *E. coli* TOP10 cells. LB agar plates supplemented with 50 μg/mL kanamycin were used for the selection of transformants at 37°C overnight. Positive clones carrying the Pr1A construct were confirmed by restriction digestion, PCR, and Sanger sequencing.

### 2.5 Construction and Cloning of the Chimeric cPr1A Gene

The Gibson assembly approach was used to construct a chimeric cPr1A gene (5′-Pr1A-BmChBD- 3′). BmChBD (*Bombyx mori* chitin-binding domain) was fused at the C-terminus of Pr1A (Protease gene of *Metarhizium anisopliae*). This required complementary overlapping sequences in the Pr1A and BmChBD segments. The Bm-Pr1A fragment, 201 bp (BmChBD with a 5′ overhang complementary to Pr1A), was amplified as described in **Section 2.6**. The Pr1A-Bm fragment, 885 bp (Pr1A with a 3′ overhang complementary to BmChBD), was amplified using Pr1A gene as template and gene-specific primers containing a vector overhang in the forward primer and BmChBD overhang in the reverse primer. Primer sequences and PCR amplification conditions given in **Supplementary Table S3** and **S4**. The amplicons were resolved by 1% gel electrophoresis and purified from the gel using the NucleoSpin Gel and PCR Clean-up kit. Subsequently, the gel-purified fragments were assembled into the purified, linearized pET-28a(+) vector using the lab- prepared Gibson assembly mix (50°C for 60 min), including the insertion of a C-terminal His-tag from the vector for affinity purification. The Gibson assembly reaction mixture (10 μL) was directly transformed into 50 μL of chemically competent *E. coli* TOP10 cells, and the transformants were selected at 37°C overnight on LB agar plates supplemented with 50 μg/mL kanamycin. Positive clones carrying the cPr1A gene insert were confirmed by restriction digestion and PCR methods.

### 2.6 Amplification of the *Bombyx mori* Chitin-Binding Domain

Primers were designed based on the DNA sequence of the *Bombyx mori* chitin-binding domain (BmChBD, NCBI accession no. AF273695.1). The amplification of the 201 bp BmChBD was done using *Bombyx mori* chitinase cDNA (clone fepM03A03, kindly provided by Prof. Toru Shimada, Gakushuin University, Tokyo, Japan) as a template and gene-specific primers containing a Pr1A overhang in the forward primer and vector overhang in reverse primer. Primer sequences and amplification conditions are provided in **Supplementary Table S3 and S5**, respectively.

### 2.7 Expression and Purification of Pr1A and cPr1A Proteins

A single colony of *E. coli* BL21(DE3) carrying the pET-28a-Pr1A or pET-28a-cPr1A constructs was inoculated into 5 mL of LB broth supplemented with kanamycin (50 µg/ mL) and incubated overnight at 37°C with shaking at 200 rpm. The overnight grown culture (1% v/v) was used to inoculate fresh LB medium, and the culture was grown at 37°C until OD_600_ reached 0.6. The protein expression was induced by adding isopropyl-1-thio-β-D-galactopyranoside (IPTG) to a final concentration of 1.0 mM, followed by incubation for 16 h at 16°C with shaking at 200 rpm. The uninduced and induced cultures were centrifuged at 13,000 rpm for 5 min at 4°C to harvest the cell pellet. Subsequently, the cell pellet was resuspended in 50 µl binding buffer (40 mM Tris- HCl, 400 mM NaCl, 10% Glycerol, 40 mM Imidazole, 0.5% Triton X-100 pH-8.0). Cells were lysed by sonication on ice using a Vibra Cell Sonicator with 4-sec pulse ON and 20-sec pulse OFF conditions.

The lysate was centrifuged at 13,000 rpm for 1 h at 4°C, and the clarified supernatant was loaded into Nickel-Nitrillo Triacetic acid (Ni-NTA) affinity chromatography column pre-equilibrated with binding buffer. Different fractions were eluted with elution buffer (40 mM Tris-HCl, 400 mM NaCl, 10% Glycerol, 400 mM Imidazole, pH-8.0). Gel imaging was done using a Bio-Rad Gel Doc EZ gel documentation system and analyzed with Image Lab software. The purified proteins were observed by SDS-PAGE and quantified by the Bradford assay.

### 2.8 Quantitative and Qualitative Analysis of Pr1A and cPr1A Recombinant Proteins

Protein concentrations of the purified recombinant Pr1A and cPr1A proteins were determined by the Lowry assay using bovine serum albumin (BSA; Sigma-Aldrich, USA) as the standard. A standard curve was generated by measuring the absorbance of BSA at concentrations ranging from 10–100 µg/mL and absorbance was measured at at 595 nm.

### 2.9 Preparation of Insect Cuticles

The Fourth-instar larvae of Eri silkworm (*Samia ricini*) were purchased from a local silk producer in Guwahati, Assam, India. The larvae were stored at −20°C overnight to freeze them for removal of the cuticle easily. The frozen larvae were gently pressed to remove the cuticles from the frozen visceral organs. The isolated cuticles were washed thoroughly under running tap water, followed by a final rinse with distilled water. The washed cuticles were dried in a hot air oven at 50°C for 5 h and then ground into a coarse powder. The cuticle powder was stored at room temperature until used as a substrate for binding and protease activity assays.

### 2.10 Comparative Cuticle-Binding Assay

A comparative cuticle-binding assay was conducted in parallel to evaluate the affinity of the Pr1A and cPr1A proteins for the *Samia ricini* cuticle under the same experimental conditions. The reaction mixture comprised of 5 mg of Eri silkworm (*Samia ricini*) cuticle powder and 2 µL (100 µg) of purified Pr1A or cPr1A enzymes suspended in 1 mL of 50 mM potassium phosphate buffer (pH 6.0). The sample mixtures were vortexed briefly to ensure uniform mixing and incubated on ice for 4 min to allow binding while minimizing protease activity. The cuticle powder was pellet down by centrifugation at 10,000 rpm for 5 min at 4°C, and the supernatant was collected. Protein concentration in the supernatant was quantified using the Lowry method. The amount of cuticle-bound protein was calculated by subtracting the unbound protein in the supernatant from the initial amount of protein added (100 µg). The relative cuticle-binding affinities of Pr1A and cPr1A were evaluated by comparing the amount (µg) of each protein bound per milligram (mg) of cuticle. All assays were performed in triplicate and the results are presented as mean ± SEM (n = 3). Relative cuticle-binding activity was obtained by normalizing the binding activity of each protein to that of Pr1A (wild type), which was set to 100%. A paired Student’s t-test was used to assess the statistical significance of the difference in binding affinity between Pr1A and cPr1A enzymes.

### 2.11 Comparative Protease Activity Assay

To compare the protease activities of Pr1A and cPr1A, the amount of peptides released from cuticular proteins during the assay was measured in parallel under identical experimental conditions. The reaction mixture containing 50 mM potassium phosphate buffer (pH 6.0), 5 mg of *Samia ricini* cuticle powder as a substrate, and 150 µg (3 µL) of purified Pr1A or cPr1A enzyme was incubated at 65°C for 4 minutes. The reaction was terminated by adding trichloroacetic acid (TCA) to a final concentration of 10% (w/v). The released peptides were quantified spectrophotometrically at 280 nm against a substrate blank as reference using L-Tyrosine as the standard curve. One unit of protease activity (U/mL) was defined as the amount of enzyme that releases 1 µg of tyrosine equivalents per minute during the specified assay conditions. The specific activity (U/mg) was calculated by normalizing the enzymatic activity against the total mass of the enzyme used in the assay. The Relative specific activity was expressed as a percentage, normalized to the specific activity of wild-type Pr1A (100%). All experiments were performed in triplicate, and the results are presented as mean ± SEM (n = 3). A paired Student’s t-test was used to determine the statistical significance between Pr1A and cPr1A specific activity.

### 2.12 Phylogenetic Analysis of Pr1A Protease

Multiple sequence alignment of various proteases was done using Clustal Omega software, and a phylogenetic tree was generated.

### 2.13 Homology Modeling and Structural Analysis of Pr1A and cPr1A Proteins

Homology modeling of Pr1A and BmChBD proteins were performed using PHYRE2 software. Secondary structures of Pr1A and cPr1A were analysed by I-TASSER. UCSF Chimera was used to align and fuse the 3D structures of Pr1A and BmChBD to generate the complete 3D structure of the chimeric cPr1A protein. The NCBI Conserved Domain Database (CDD) was used to identify the conserved domains.

### 2.14 Statistical Analysis

All experiments were performed in independent triplicates. Data analysis was done using a paired Student’s t-test with a p value of *p* < 0.05 considered statistically significant.

**Image 1.**
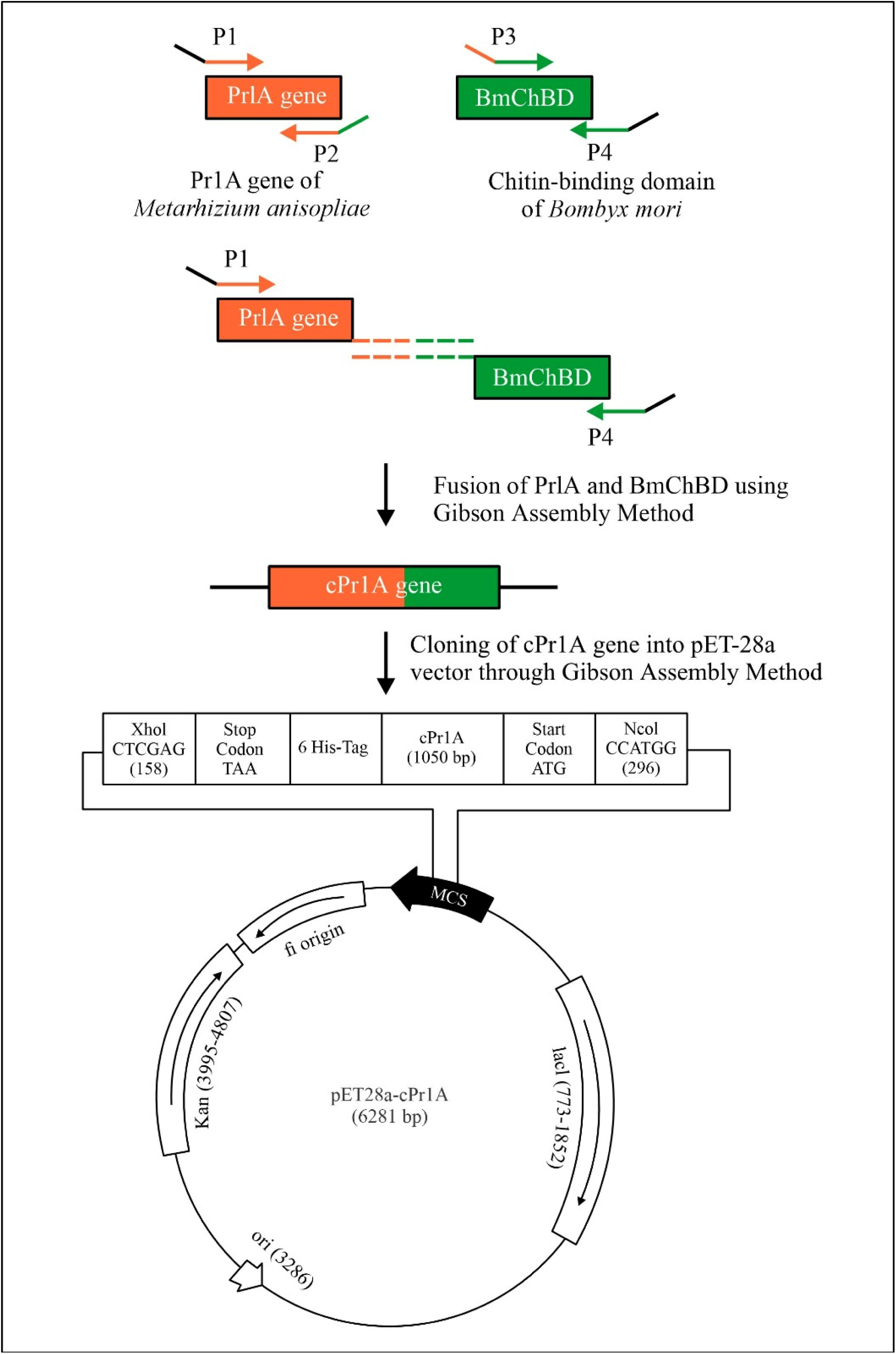
Pictorial illustration of construction and cloning of chimeric cPr1A gene into pET-28a

## 3. Results

### 3.1 Cloning of Wild-Type Pr1A and Chimeric cPr1A Genes

The wild-type Pr1A protease gene was amplified from the cDNA of *Metarhizium anisopliae* yielding a 906 bp amplicon band corresponding to the Pr1A gene **(Figure 1)**. The gel-purified amplicon was cloned into the NcoI/XhoI digested pET-28a(+) vector using the Gibson assembly, resulting in pET-28a-Pr1A (6145 bp).

**Figure 1.**
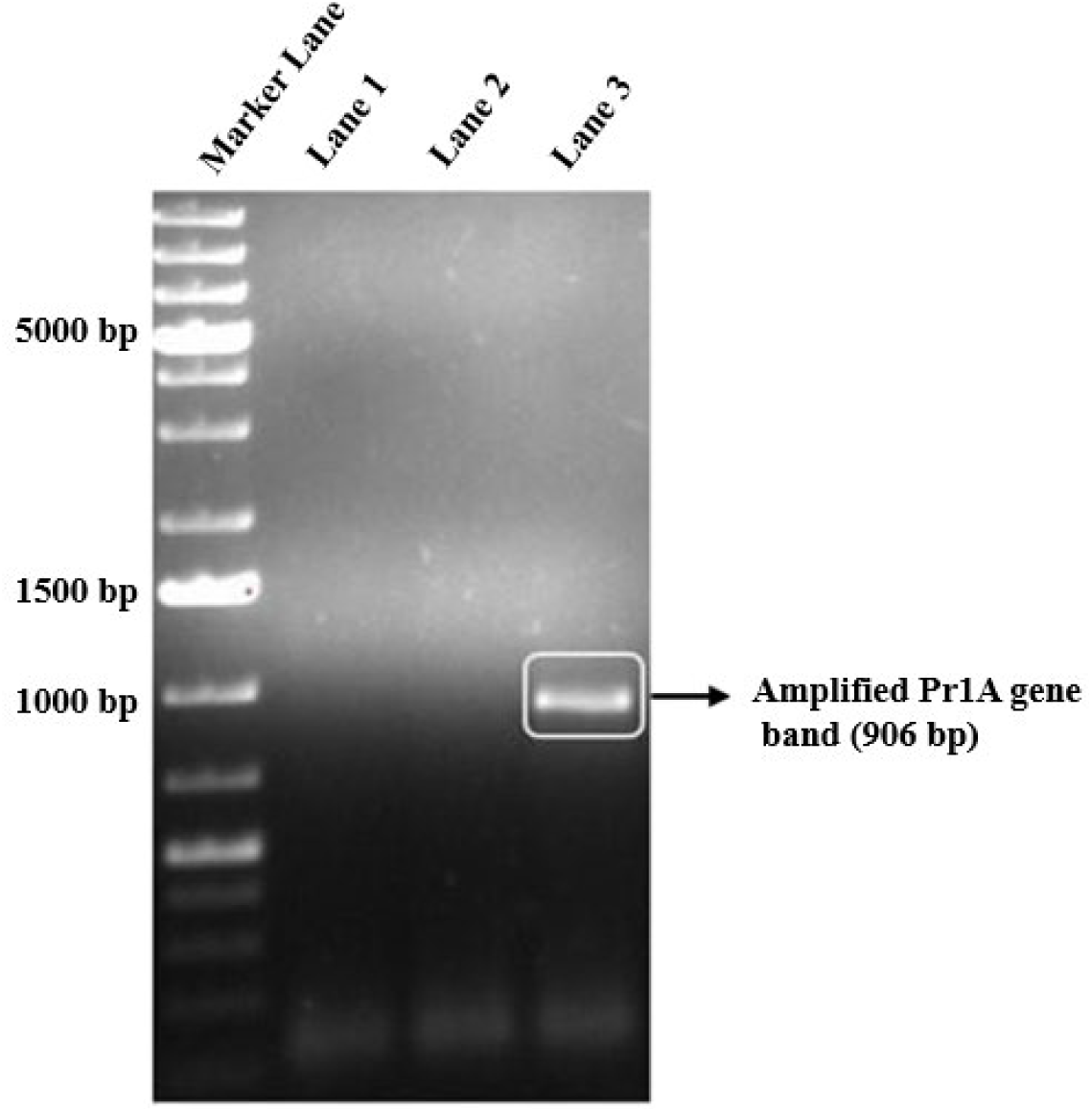
Agarose gel electrophoresis showing amplified Pr1A gene from cDNA of *Metarhizium anisopliae* using gene-specific primers. PCR was done at annealing temperatures, 64°C (Lane 1), 66°C (Lane 2), 68°C (Lane 3) to optimize the annealing temperature. Marker Lane: 1 Kb plus DNA Ladder (Thermo Scientific). Lane 3: Amplified Pr1A gene band (906 bp)

The chimeric cPr1A chimeric gene was assembled by fusing the two overlapping fragments: Pr1A- Bm, (885 bp) and Bm-Pr1A (201 bp). Both fragments were amplified and verified by agarose gel electrophoresis **(Figure 2)**. The gel-purified Pr1A-Bm and Bm-Pr1A fragments were cloned into the NcoI/XhoI digested pET-28a(+) vector to produce the recombinant plasmid pET-28a-cPr1A (6281 bp) containing the full-length chimeric cPr1A gene (1050 bp). The recombinant plasmids pET-28a-Pr1A (6.1 kb) and pET-28a-cPr1A (6.3 kb) were transformed and positive clones were screened by restriction digestion and the PCR method. The in-frame fusion between Pr1A & pET- 28a(+), and cPr1A & pET28a vector at the Gibson junction was confirmed by Sanger sequencing of positive clones without any mutation. Sanger sequencing of the mature Pr1A DNA sequence identified two nucleotide variations when compared to the reference sequence from NCBI (accession No. M73795.1), occurring at nucleotide positions [650] and [664] in the complete sequence (after exclusion of the signal and propeptide regions). These variations resulted in amino acid changes from [Valine → Glycine] at the mature residue [217] and from [Serine → Threonine] at the mature residue [222] (with the mature domain numbering starting from position 1 at the N-terminal catalytic residue). **(Supplementary Figure S1 and S2)**.

**Figure 2.**
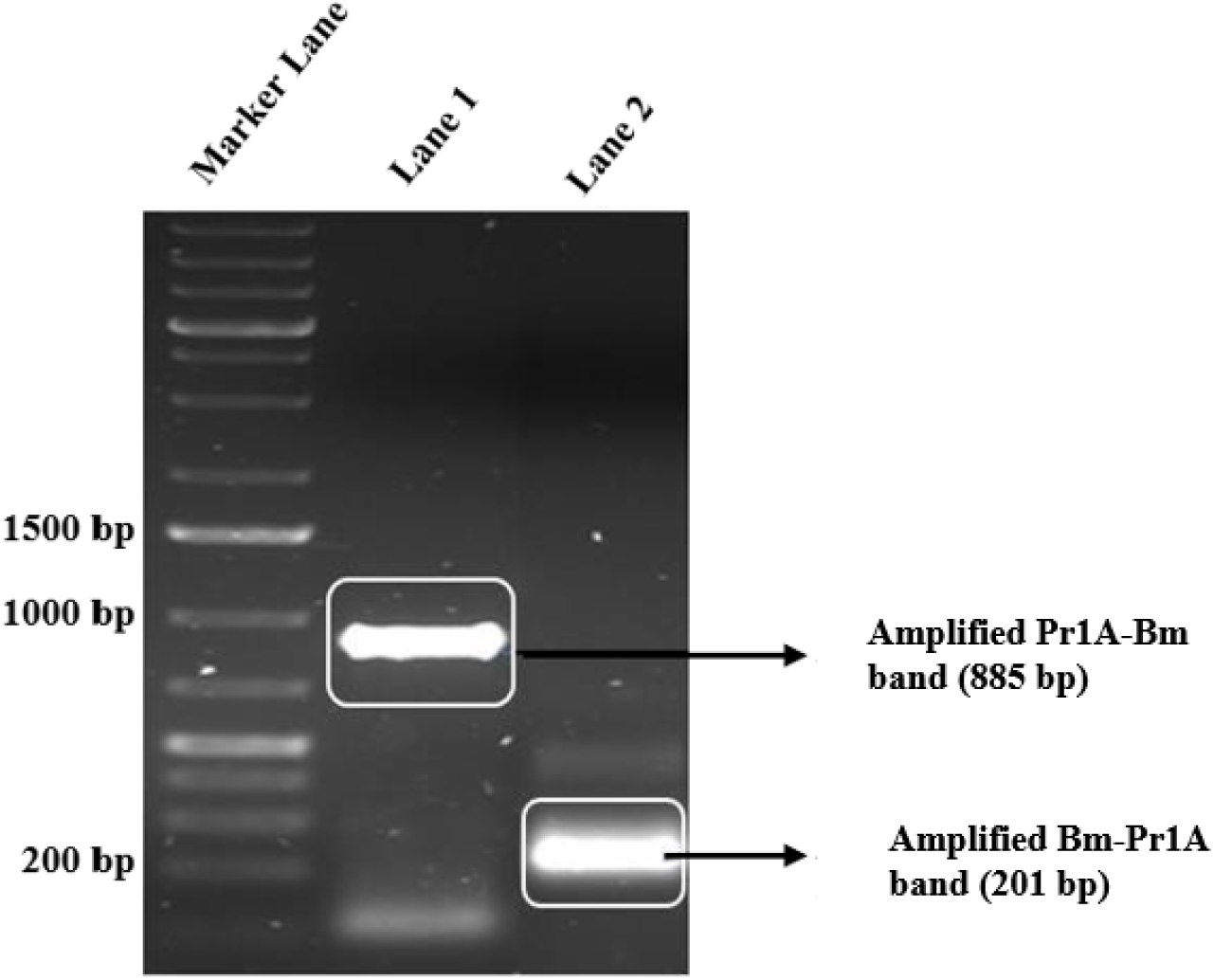
Agarose gel electrophoresis image showing fragments of Pr1A-Bm and Bm-Pr1A with overlapping ends for construction of the chimeric cPr1A gene. A 1% agarose gel was used to resolve the PCR products Marker lane: 1 Kb plus DNA Ladder (Thermo Scientific). Lane 1: Pr1A-Bm fragment (885 bp) – Pr1A gene with *Bombyx mori* chitin-binding domain overhang at 3’ end. Lane 2: Bm-Pr1A fragment (201 bp) – *Bombyx mori* chitin-binding domain with Pr1A overhang at 5’ end.

### 3.2 Confirmation of Pr1A & cPr1A Positive Clones

To confirm the cloning of wild-type Pr1A and chimeric protease cPr1A, the plasmids extracted from the Pr1A and cPr1A transformant colonies were digested with NcoI and XhoI restriction enzymes, and the resulting digested fragments were visualized on agarose gel.

The agarose gel analysis confirmed the presence of cloned gene inserts in the 4th, 9th, 11th, and 12th (out of 15 screened) Pr1A colonies and the 1st to 8th (out of 13 screened) cPr1A colonies and revealed four DNA bands corresponding to the linearized pET-28a vector (5369 bp) and the respective inserts: Pr1A, 906 bp (217 bp, 467 bp, and 185 bp), and cPr1A, 1050 bp (217 bp, 467 bp, and 329 bp) **(Figure 3 and Figure 4)**, which confirmed the successful cloning of both Pr1A and cPr1A inserts into the pET-28a vector. The expected single 906 bp and 1050 bp fragments were instead observed as three distinct bands, since there are two NcoI restriction sites within the Pr1A gene (at 236 bp and 703 bp). The increased size of the third fragment of cPr1A (329 bp) is consistent with the fusion of the BmChBD domain at the C-terminus of Pr1A. Two minor fragments (18 bp and 19 bp), originating from vector-insert junctions, were anticipated but were too small to detect on the gel. Adding the observed bands to these small fragments (a combined 37 bp) results in estimated total insert sizes of 906 bp (Pr1A) and 1050 bp (cPr1A), aligning with the expected chimeric construct. The two nucleotide variations identified in the mature Pr1A sequence (Val217Gly and Ser222Thr) might be due to strain variations of *Metarhizium anisopliae*.

**Figure 3.**
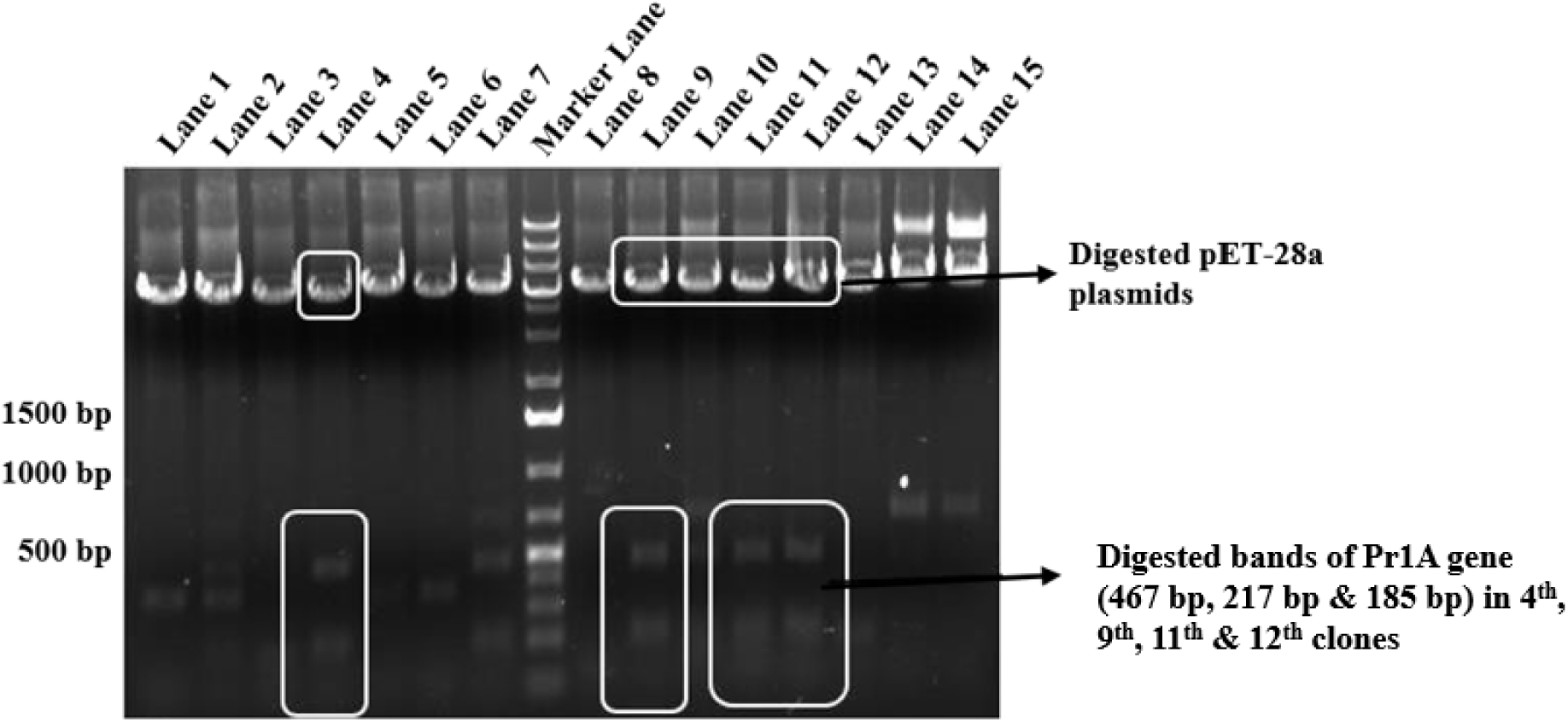
Agarose gel electrophoresis showing restriction digestion analysis for confirmation of positive (pET-28a - Pr1A) clones. Plasmids were digested with NcoI and XhoI restriction enzymes. Marker lane – 1 Kb plus DNA Ladder (Thermo Scientific). Lane 4, 9, 11 & 12: Positive clones showing release of the Pr1A gene insert. The Pr1A gene has two internal NcoI restriction sites at 213 bp and 703 bp therefore released three bands 467 bp, 217 bp, and 185 bp instead of a single 906 bp band of Pr1A.

**Figure 4.**
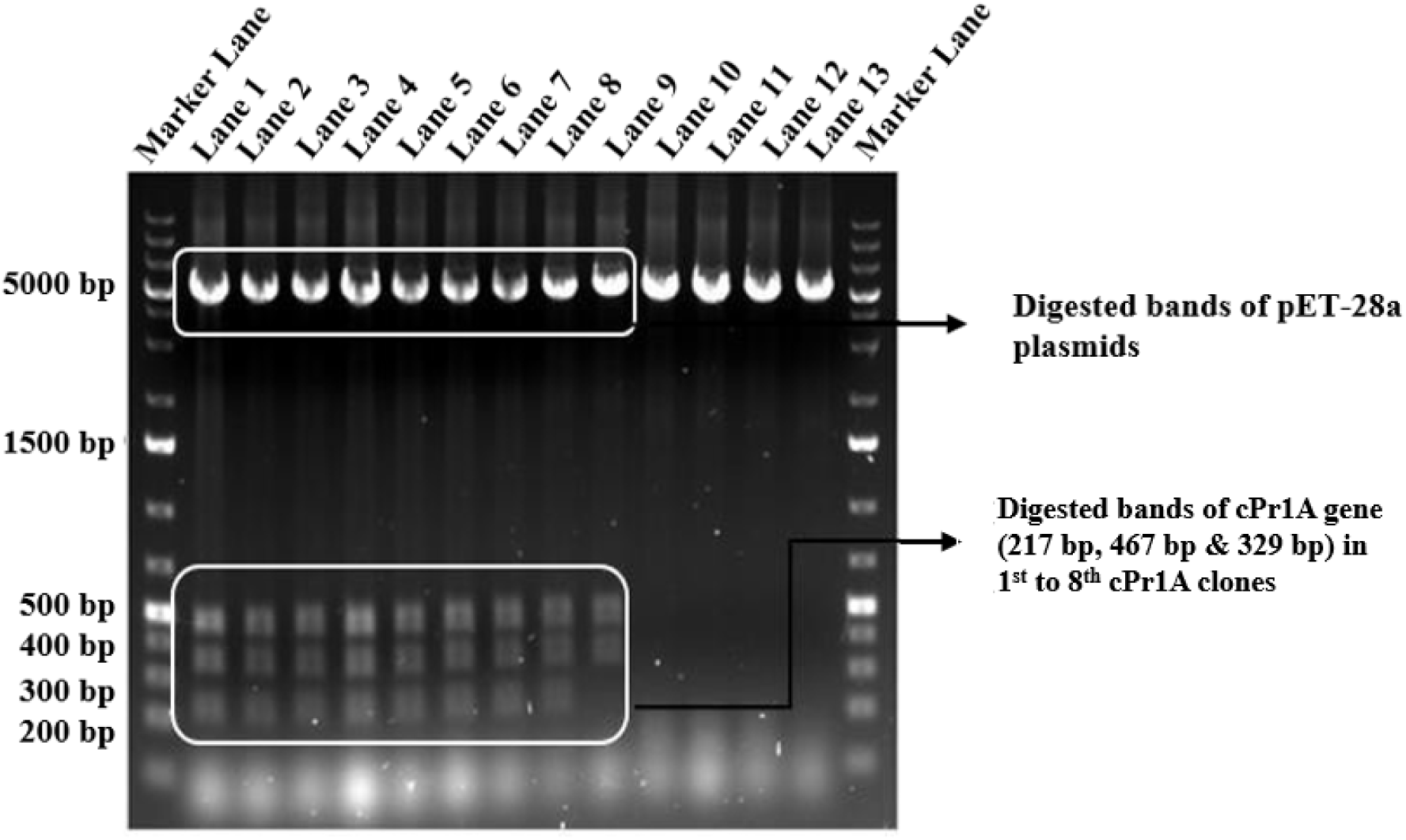
Agarose gel electrophoresis showing restriction digestion analysis for confirmation of positive pET-28a - cPr1A clones. Plasmids were digested with NcoI and XhoI restriction enzymes. Marker lane: 1 Kb plus DNA Ladder (Thermo Scientific). Lane 1 to 8: Positive clones showing release of the cPr1A gene insert. The cPr1A gene has two NcoI restriction sites at 213 bp and 703 bp, giving three bands 467 bp, 217 bp, and 329 bp instead of a single 1050 bp band.

PCR amplification employing gene-specific primers provided a second confirmation of positive clones. Agarose gel analysis of PCR products indicated that the 1st to 12th Pr1A plasmids (out of 12 screened) and the 1st to 8th cPr1A plasmids (out of 8 screened) carried the respective Pr1A and cPr1A genes **(Figure 5 and Figure 6)**. Positive colonies were subsequently grown in 10 mL LB media, and plasmids were isolated and stored at -20°C for protein expression experiments.

**Figure 5.**
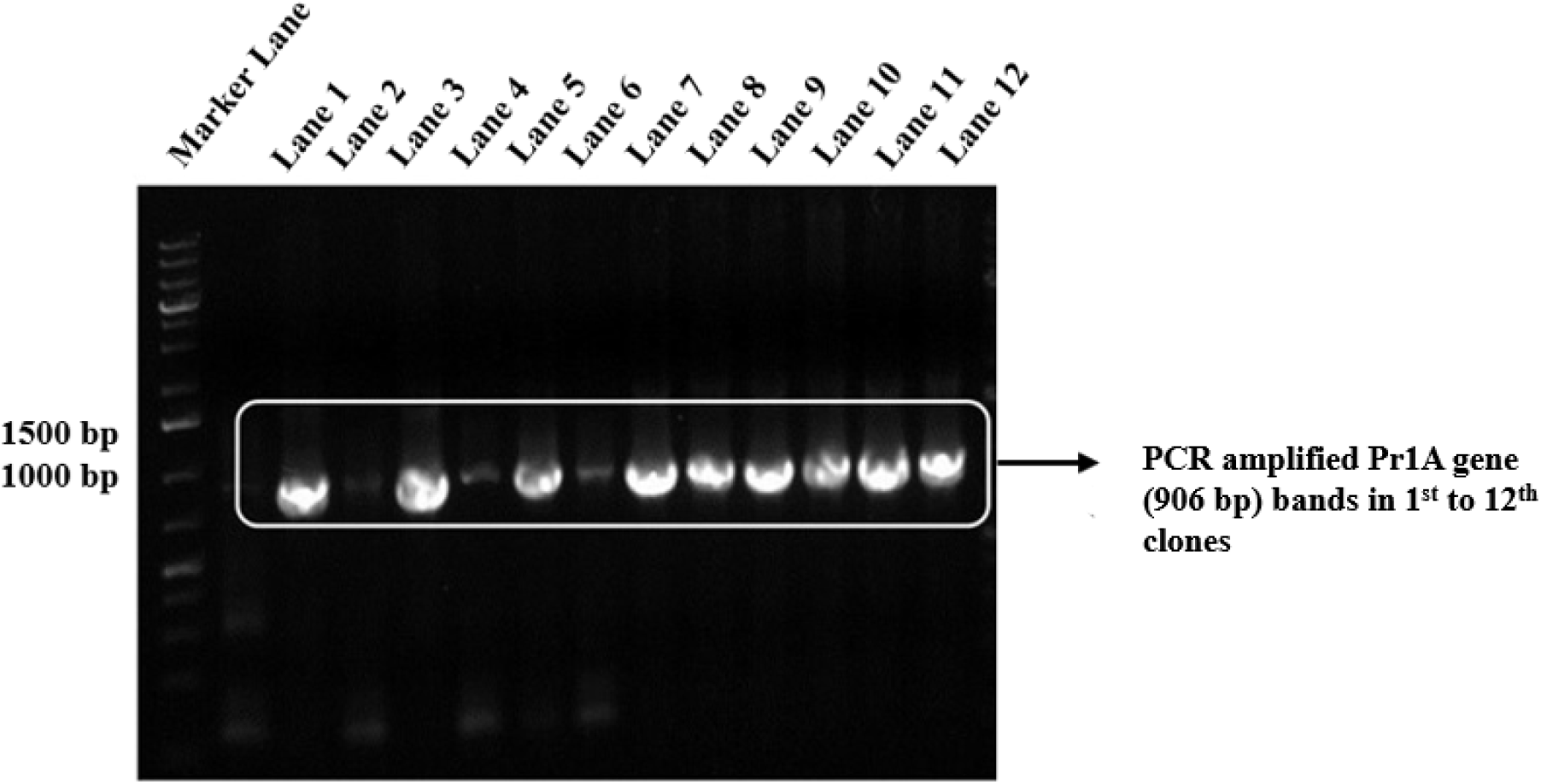
Agarose gel analysis showing PCR amplified Pr1A gene (906 bp) bands from Pr1A clones. PCR confirmation for positive clones was done using Taq DNA polymerase and Pr1A gene-specific primers with plasmid DNA isolated from cPr1A clones 1st to 12th as template Marker lane: 1 Kb plus DNA Ladder (Thermo Scientific). Lane 1 to 12: Clones 1st to 12th showing amplified Pr1A gene insert (906 bp).

**Figure 6.**
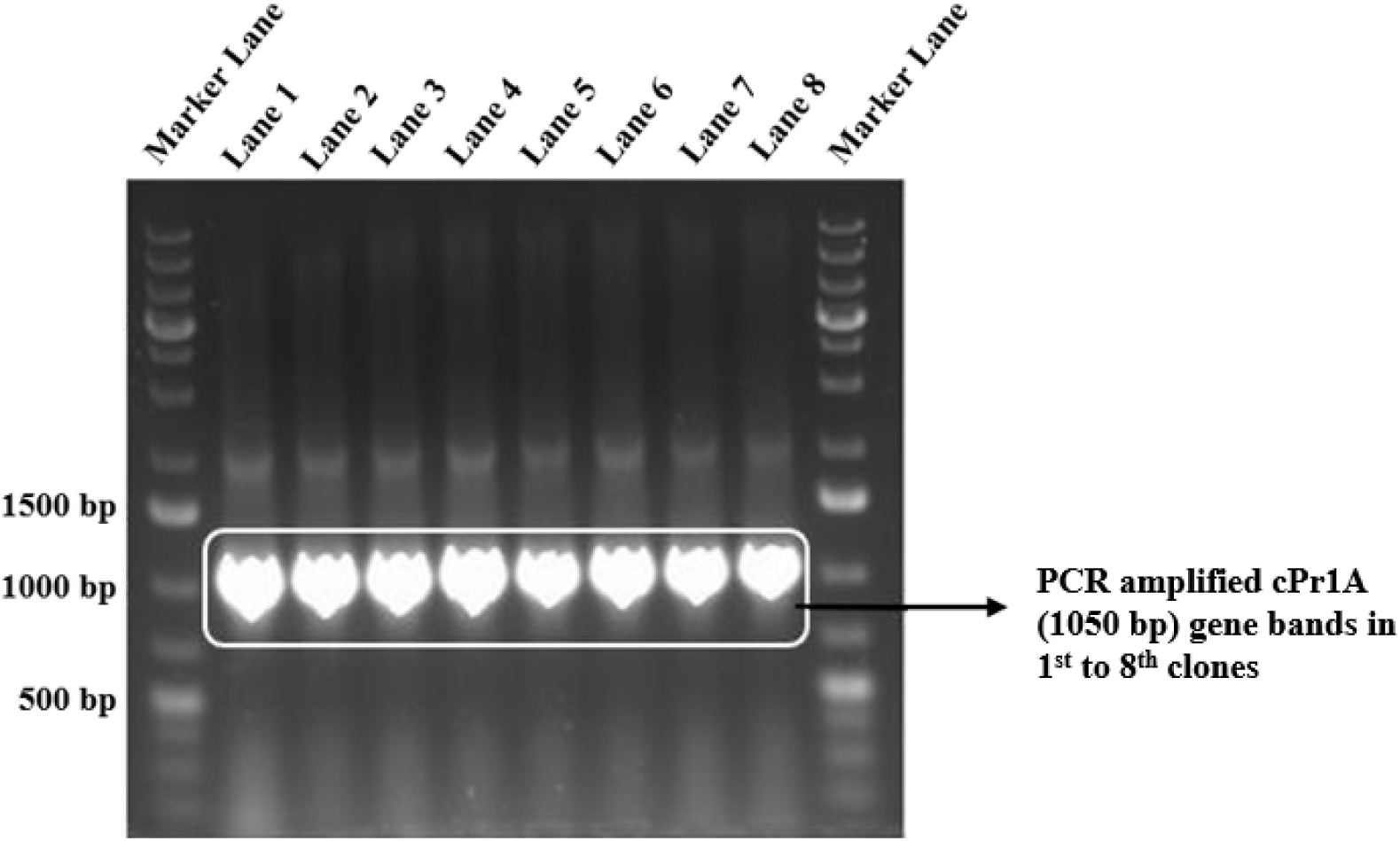
Agarose gel analysis showing PCR amplified cPr1A gene (1050 bp) fragments from cPr1A clones. PCR confirmation for positive clones was done using Taq DNA polymerase and cPr1A gene-specific primers with plasmid DNA isolated from cPr1A clones 1st to 8th as templates. Marker lane – 1 Kb plus DNA Ladder (Thermo Scientific). Lane 1 to 8 – Clones 1st to 8th showing amplified cPr1A gene insert (1050 bp).

### 3.3 Expression and Affinity Purification of Wild-Type (Pr1A) and Chimeric (cPr1A) Proteins

The recombinant plasmids harbouring the Pr1A and cPr1A genes were transformed into *E. coli* BL21(DE3) competent cells. Both Pr1A-(His)_6_ and cPr1A-(His)_6_ proteins were overexpressed in LB broth. Pr1A and cPr1A proteins were expressed as soluble proteins, with purified molecular weights of approximately 29.4 kDa (Pr1A) and 34.9 kDa (cPr1A) on SDS-PAGE, consistent with theoretical predictions. The purification of wild-type Pr1A and chimeric cPr1A proteins were done using affinity chromatography. Protein concentrations were determined using the Lowry method with Bovine Serum Albumin (BSA) as a standard. Purified proteins, along with eluted fractions from the column were analysed on SDS-PAGE **(Figure 7 and Figure 8).** The amount of wild-type Pr1A and chimeric cPr1A proteins obtained from the one litre culture was 87 mg and 93 mg, respectively.

**Figure 7.**
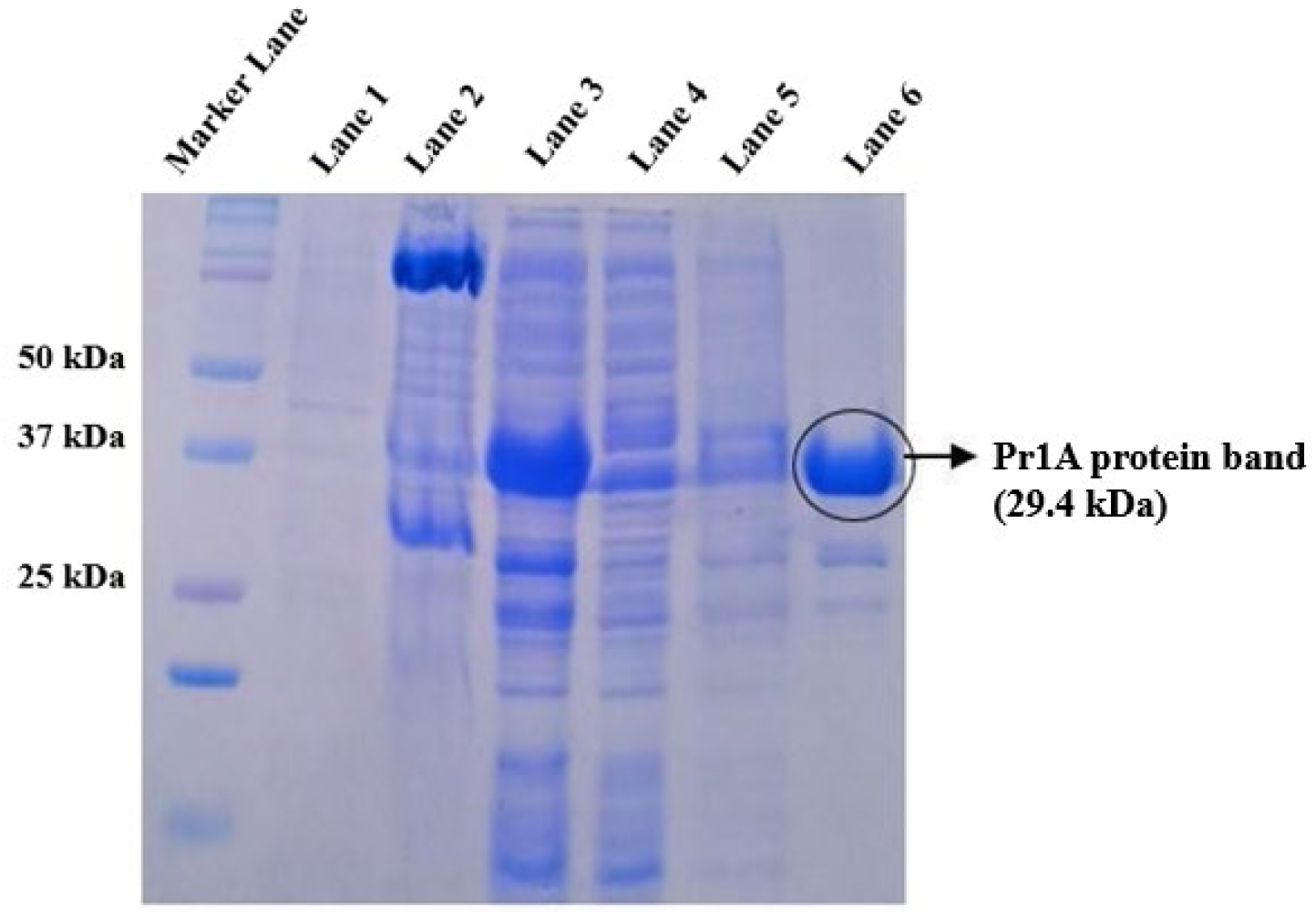
Purification profile of recombinant Pr1A protein. Proteins were resolved by 12% (w/v) SDS-PAGE and stained with Coomassie Brilliant Blue. Marker Lane: Bio-Rad Precision Plus Protein Dual Color Ladder (Catalogue #1610374). Lane 1: Cell pellet after sonication. Lane 2: Soluble Supernatant after sonication. Lane 3: Flow-through fraction. Lane 4: Last column wash. Lane 5: Purified Pr1A protein.

**Figure 8.**
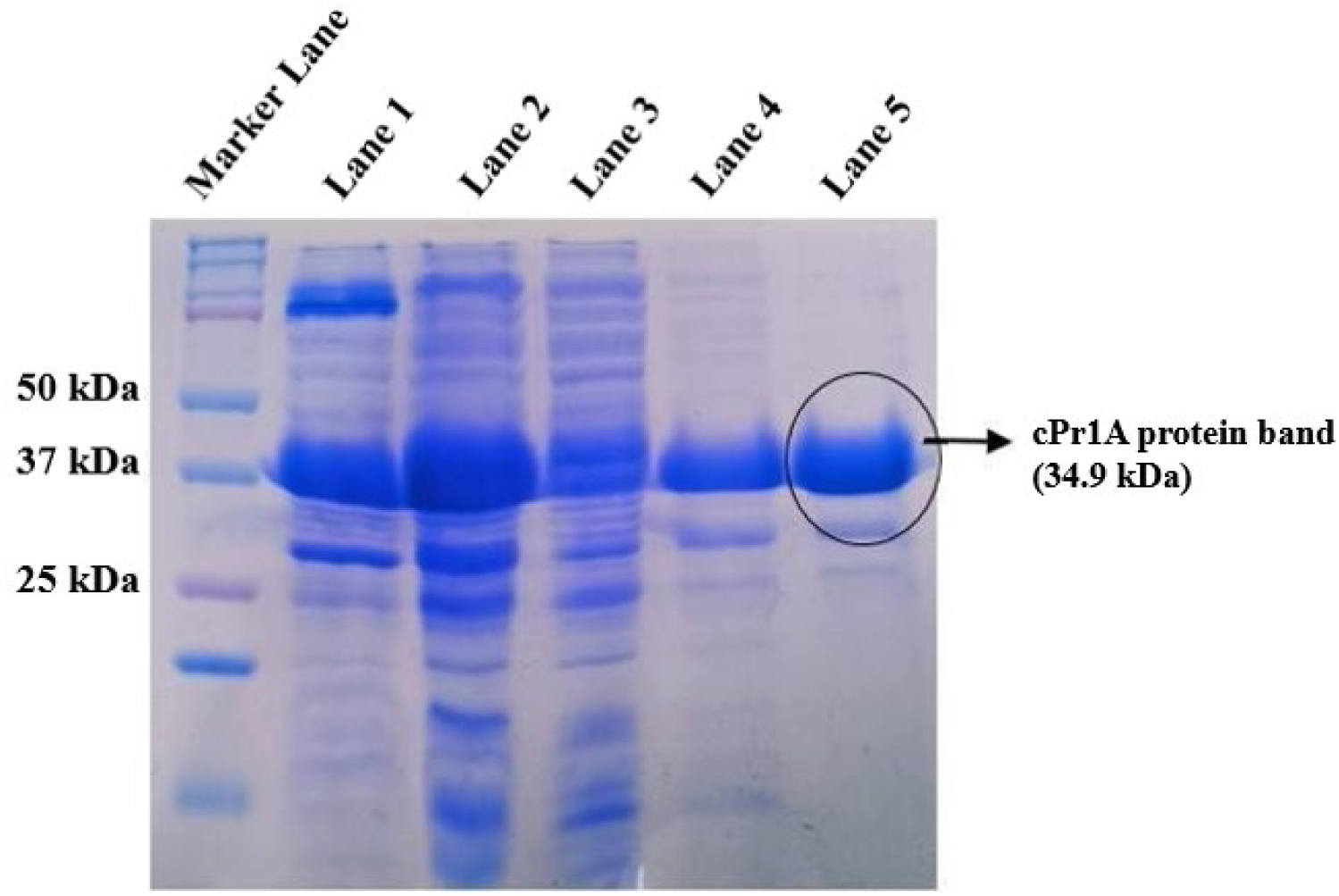
Purification profile of recombinant cPr1A protein. Proteins were resolved by 12% (w/v) SDS-PAGE and stained with Coomassie Brilliant Blue. Marker Lane: Bio-Rad Precision Plus Protein Dual Color Ladder (Catalogue #1610374). Lane 1: Cell pellet after sonication. Lane 2: Soluble Supernatant after sonication. Lane 3: Flow-through fraction. Lane 4: Last column wash. Lane 5: Purified cPr1A protein.

### 3.4 Comparative Cuticle-Binding Assay

The wild-type Pr1A and chimeric protease cPr1A were successfully expressed in *E. coli* and purified by Ni-NTA affinity chromatography. Binding assays revealed significantly enhanced binding affinity of the chimeric (cPr1A) compared to wild-type Pr1A towards the cuticle of *Samia ricini* silkworm. When incubated under identical conditions, the mean amount of bound protein was 12.27 ± 2.13 µg/mg cuticle powder for wild-type Pr1A and 15.81± 1.97 µg/mg cuticle powder for chimeric cPr1A (n=3 independent replicates, mean ± SEM), corresponding to an approximately 28.9% higher binding for the chimeric protease **(Figure 9)**. A paired Student’s t-test confirmed a statistically significant difference (*p* < 0.002).

**Figure 9.**
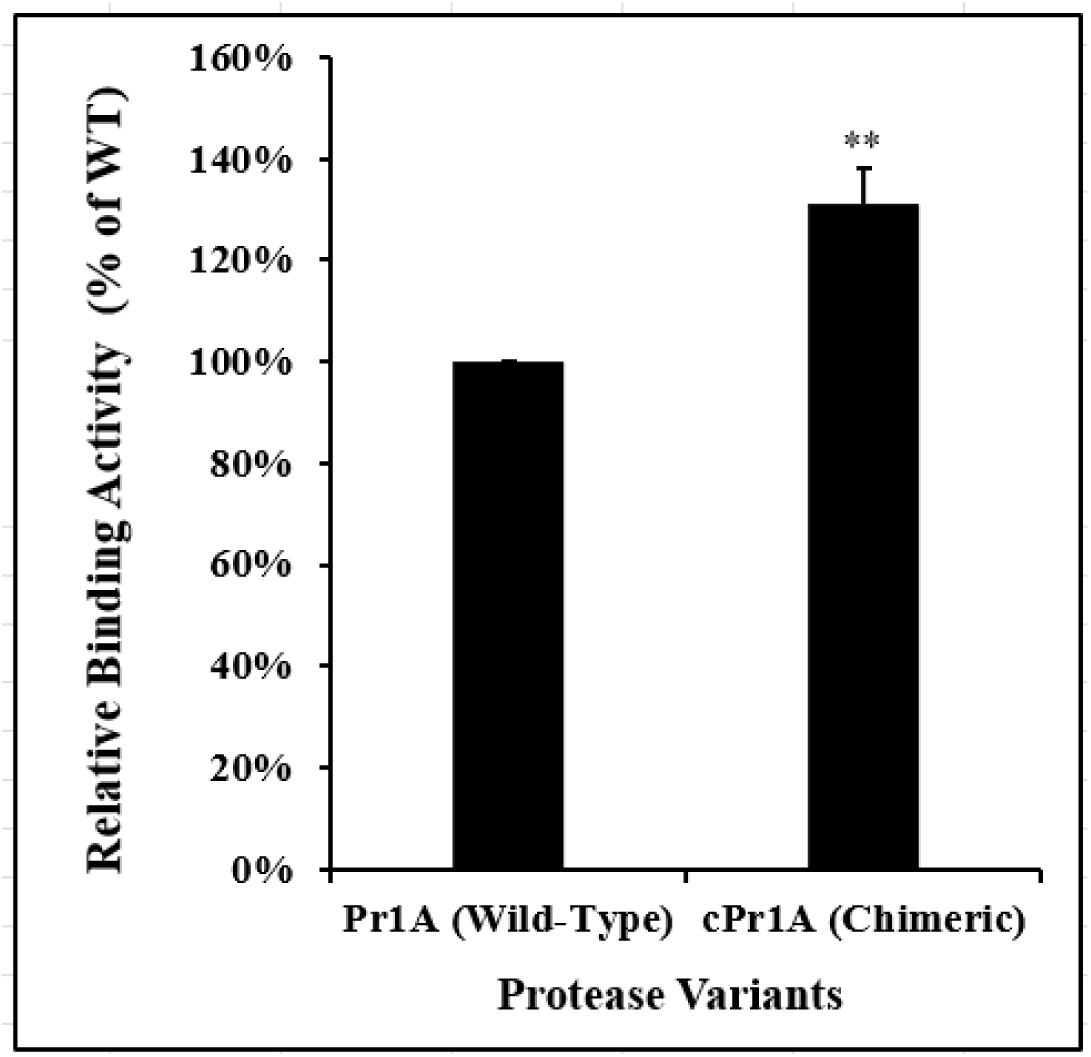
Comparative cuticle binding analysis of wild-type Pr1A and chimeric cPr1A protease. The binding assay was performed using 5 mg of *Samia ricini* cuticle powder in 50 mM potassium phosphate buffer (pH 6.0) containing 100 μg of protease. After incubation, the amount of unbound protein in the supernatant was quantified, and the bound protein was calculated by subtraction from the initial 100 μg. Values represent mean ± standard deviation from three independent experiments. Pr1A (wild-type): 12.27 ± 2.13 μg bound protein per mg cuticle powder cPr1A (chimeric): 15.81± 1.97 μg bound protein per mg cuticle powder The chimeric protease cPr1A demonstrated a significantly 28.9% higher binding affinity compared to the wild-type Pr1A. This difference was statistically significant (paired t-test, *p* < 0.002).

### 3.5 Comparative Protease Activity Assay

To assess whether fusion of the *Bombyx mori* chitin-binding domain BmChBD to the Pr1A could improve the protease activity of chimeric cPr1A compared to wild-type Pr1A, we conducted protease assay experiments and measured the specific activities of both proteins. WT Pr1A showed a specific activity of 0.254 ± 0.06 U/mg protein, while cPr1A reached 0.343 ± 0.08 U/mg protein (n=3 independent replicates, mean ± SEM) **(Supplementary Table S6)**. The chimeric cPr1A exhibited significantly higher specific activity compared to wild-type Pr1A (35% increase, *p* < 0.03; paired Student’s t-test) this increased specific activity of chimeric protease was consistent with the enhanced binding affinity observed in the comparative cuticle binding assay. cPr1A showed a relative specific activity of 135% when normalised to WT Pr1A set at 100% **(Figure 10)**.

**Figure 10.**
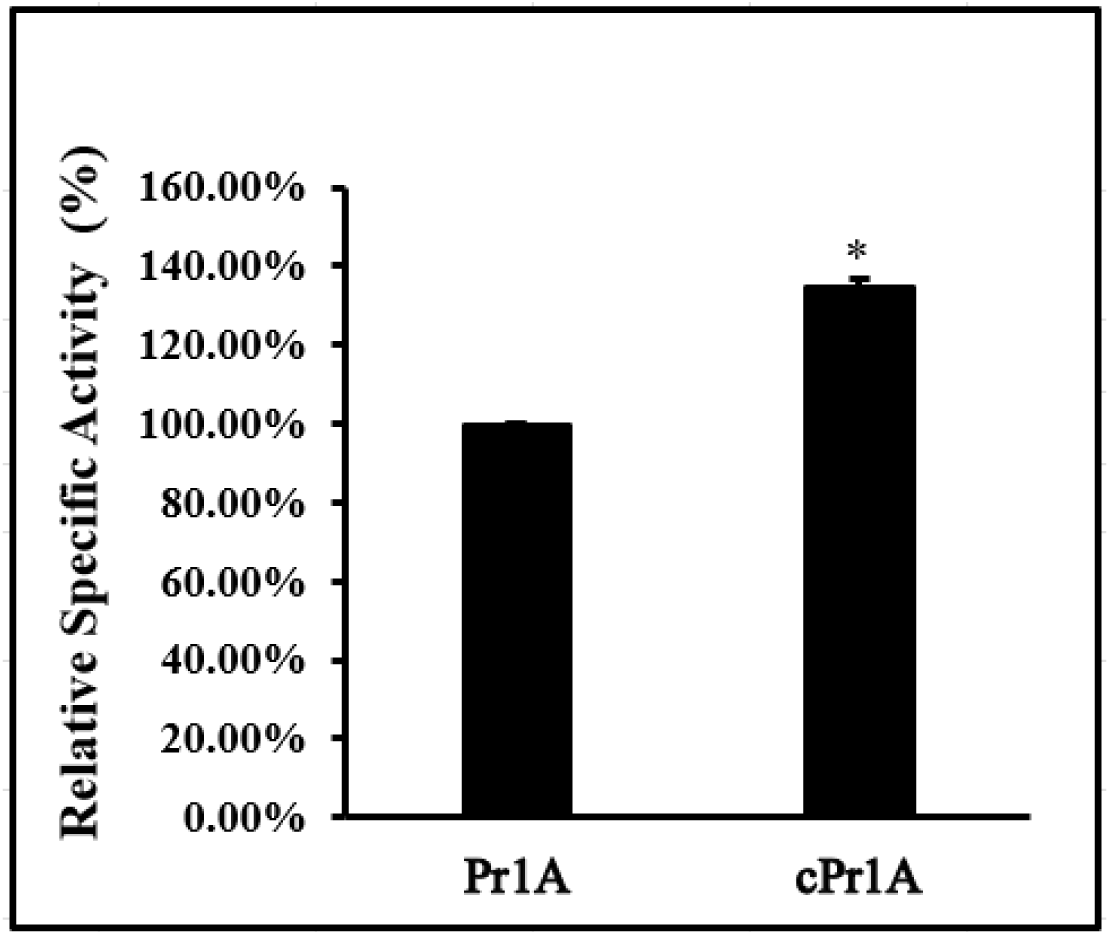
Comparative enzymatic analysis of purified wild-type Pr1A protease and chimeric cPr1A protease using *Samia ricini* cuticle powder as substrate. Relative specific activity is expressed as a percentage of wild-type Pr1A (set as 100%). The chimeric protease cPr1A exhibited significantly 35% higher enzyme activity. Absolute specific activity values: Wild-type Pr1A = 0.254 ± 0.06 U/mg. Chimeric cPr1A = 0.343 ± 0.08 U/mg. Statistical significance was determined by a paired two-sample t-test (*p* < 0.03) using absolute activity values. Values represent the means of three independent experiments.

### 3.6 Homology Modeling and Structural Analysis of Pr1A and cPr1A Proteins

Homology modeling of Pr1A and BmChBD proteins showed that WT Pr1A shared 100% sequence coverage with cuticle-degrading protease from *Paecilomyces lilacinus* (PDB: 3F7O) with 100% confidence while BmChBD revealed 99.4% confidence and 96% coverage with avirulence protein 4 (avr4) from *Cladosporium fulvum*. The 3D structure of Pr1A is shown in **(Figure 11)**. The secondary structure analysis showed a difference between WT Pr1A and cPr1A. WT Pr1A consisted of 56.6% random coils, 17.99% β-strands, and 25.4% helix, while cPr1A consisted of 59.9% random coils, 18.34% β-strands, and 21.7% helix **(Supplementary Table S7)**. The 3D protein structure of cPr1A was generated by fusing the 3D structures of Pr1A and BmChBD using UCSF Chimera software **(Figure 12)**. The conserved domain analysis of *M. anisopliae* protease identified two highly conserved domains: Peptidases S8 (PCSK9/proprotein convertase subtilisin/kexin type 9 & Proteinase K) and the Peptidase S53 superfamily.

**Figure 11.**
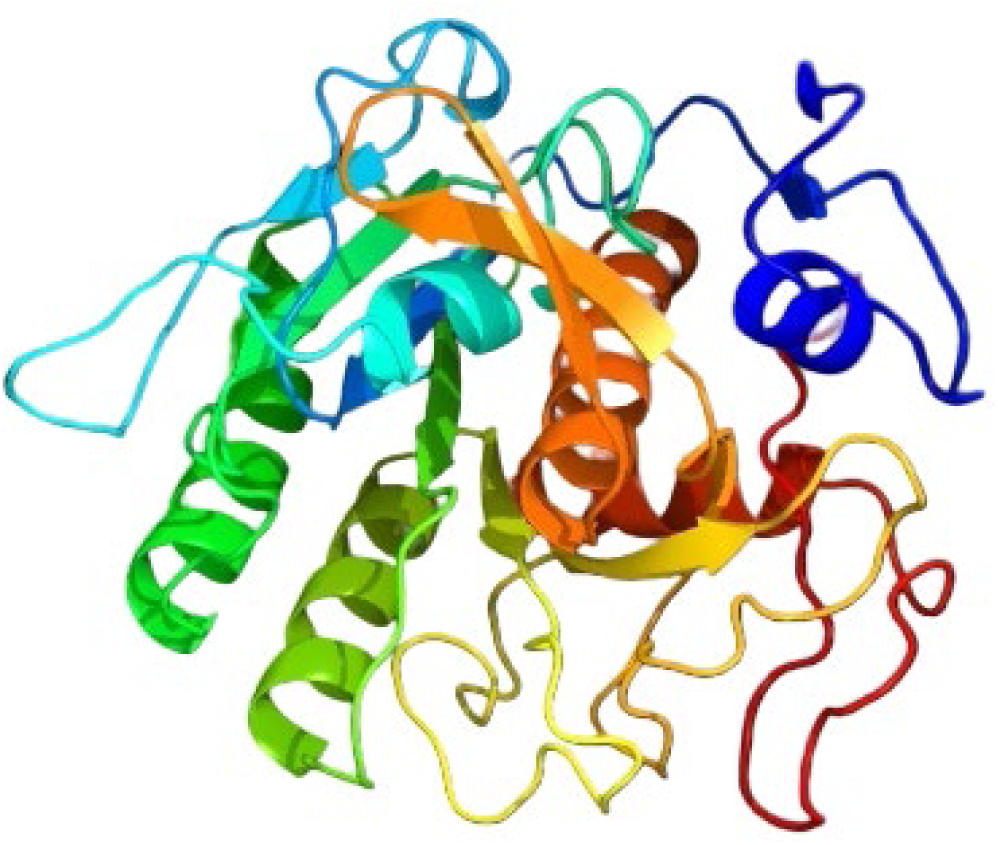
3D protein structure of wild-type Pr1A protease was modelled using PHYRE2 web server. The model exhibits 100% confidence and 100% sequence coverage. The model shares 76% identity with the template structure PDBe-3f7o (Crystal structure of Cuticle-Degrading Protease from *Paecilomyces lilacinus*, PL646). Predicted structure is shown in Ribbon representation.

**Figure 12.**
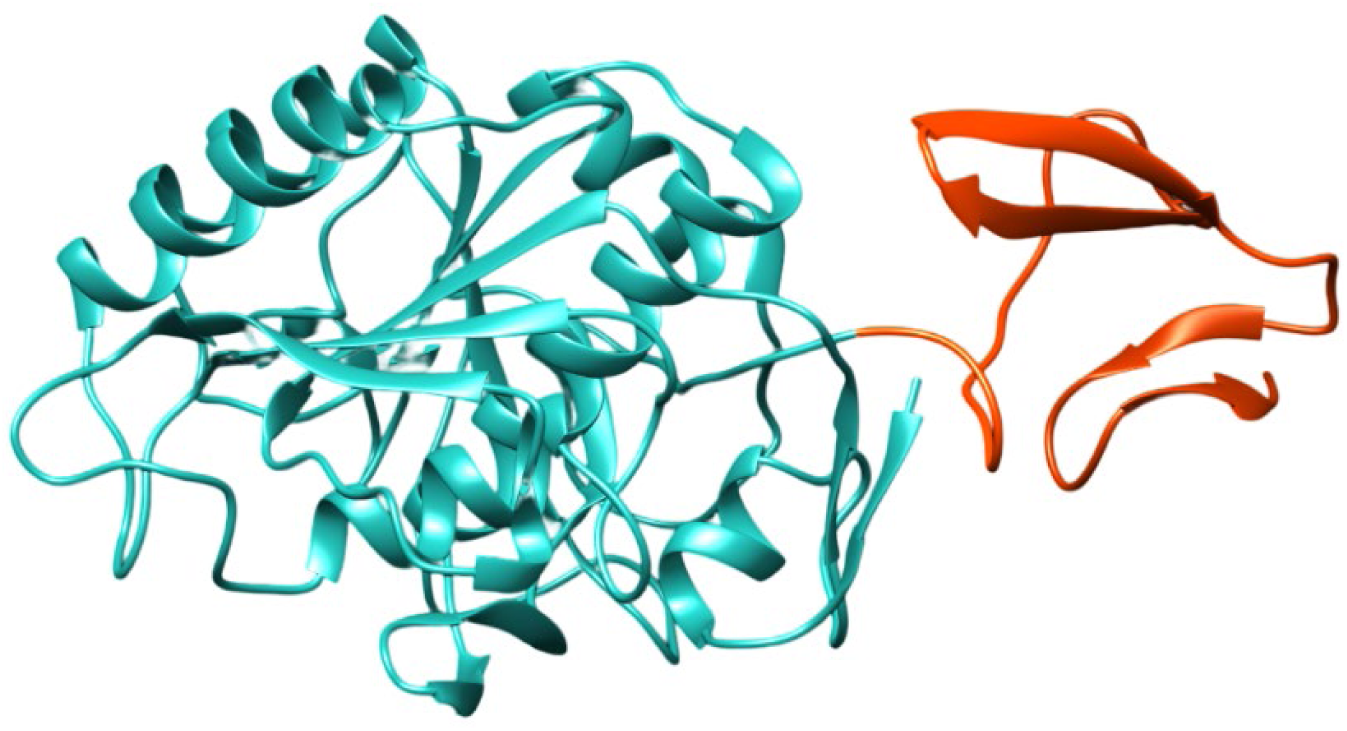
3D protein structure model of chimeric cPr1A protease. The structure of BmChBD (*Bombyx mori* chitin-binding domain) was generated using the PHYRE2 web server with 99.4% confidence and 96% sequence coverage, sharing 35% identity with the template structure PDBe-6bn0 (Avirulence protein 4, Avr4, from *Cladosporium fulvum* bound to chitin hexasaccharide). The chimeric cPr1A protease was constructed by structural fusion of the modelled Pr1A and BmChBD domains using UCSF Chimera software. The Pr1A domain is depicted in green; the BmChBD domain is depicted in red. The ribbon representation shows the overall architecture and domain orientation of the chimeric construct.

### 3.7 Phylogenetic Analysis of Pr1A Protease

Multiple sequence alignment was performed using Clustal Omega, and a phylogenetic tree was generated from the alignment using MEGA 11 software **(Figure 13)**. The sequence homology of the *M. anisopliae* protease gene with protease genes reported from other organisms was investigated using the BLASTn (nucleotide) and BLASTp (protein) programs. At the protein level, *M. anisopliae* Pr1A protease showed 76% identity with subtilisin Pr1A from *Paecilomyces lilacinus*, 48% identity with *Thermus aquaticus*. At the nucleotide level, the Pr1A protease of *M. anisopliae* exhibited 68% identity with the alkaline protease of *Parengydontium album sp.* and 82% sequence identity with the P32 gene from the *Metapochonia rubescens* strain CBS. The Pr1A protease gene also showed sequence identity with the genes from other organisms, including *Lecanicillium psalliotae* (70%) and *Vibrio sp.* PA-44 (45%), *Lederbergia lenta* (35%), *Thermococcus kodakarensis* (32%), *Thermoactinomyces vulgaris* (39%), and *Homo sapiens* (35%).

**Figure 13.**
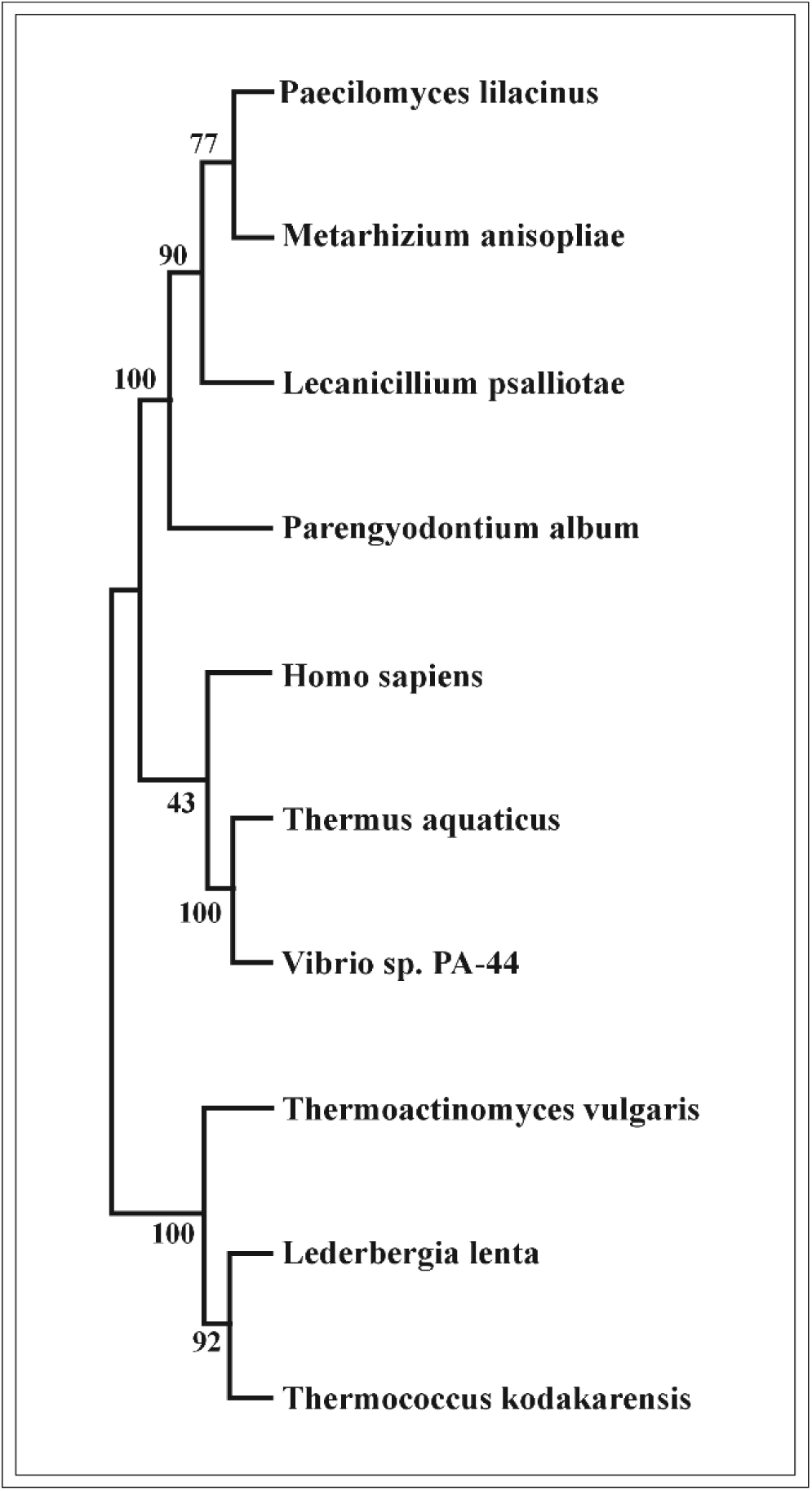
Phylogenetic tree depicting the evolutionary relationship of *M. anisopliae* Pr1A protease with proteases from related organisms. The phylogenetic tree was constructed using the MEGA version 11 software, employing the neighbour-joining approach with the Poisson model for multiple amino acid substitutions and branches. 1000 random bootstrap repeats. Sequence alignment was done using Clustal Omega. Binomial genus-species names were used. High bootstrap values (close to 100%) indicate strong support for the branches. The evolutionary relationship revealed that *Metarhizium anisopliae’s* Pr1A protease was most closely related to the Pr1A protease of *Paecilomyces lilacinus*.

### 3.8 Discussion

The entomopathogenic fungus *Metarhizium anisopliae* is an essential part of sustainable pest management strategies because it can naturally infect and kill insect hosts through the synergistic action of enzymatic cuticle degradation and mechanical pressure from appressoria. Extracellular enzymes, particularly subtilisin-like proteases and chitinases, are key virulence determinants of the fungi that facilitate host cuticle penetration and nutrient acquisition for the fungi [20–24]. Among the array of 11 secreted proteases produced by *Metarhizium anisopliae* during the pathogenesis, Pr1A, a subtilisin-like protease, and Pr4, a cysteine protease, are identified as the key enzymes primarily involved in the degradation of insect cuticles in the early penetration stage [25–26]. Pr1A has been the major target for genetic modification to enhance the entomopathogenicity of the fungus. One approach involves the fusion of chitin-binding domains (ChBDs) to these protease enzymes. A chitin-binding domain ChBD identified in certain chitinases, including those found in plants and insects, plays a pivotal role in the efficient binding of the enzyme to crystalline chitin substrates by positioning the catalytic domain in proximity to the substrate, increasing enzyme concentration on the substrate, and enhancing the enzyme activity towards crystalline chitin. The critical role of ChBDs for the functional properties of the chitinases is confirmed by their deletion, which resulted in a significant reduction in the catalytic efficiency and the binding affinity for crystalline chitin substrates [27–35].

Many proteases containing chitin-binding domains have been found in bacteria and insects, such as Sp22D from *Anopheles gambiae* and AprIV from *Alteromonas sp.* [36–38]. Furthermore, Protease C with a C-terminal chitin-binding domain is found in *Streptomyces griseus*, suggesting that domain fusion is a conserved strategy to improve substrate interaction and support chitin-mediated processes like cuticle digestion and host defense mechanisms [28–29, 39]. Fan et al. demonstrated that fusing a *Bombyx mori* chitin-binding domain (BmChBD) to the C-terminal of subtilisin-like protease CDEP-1 from *Beauveria bassiana* boosts its binding affinity to the cuticular chitin and increases the hydrolysis of cuticular proteins, accelerates penetration, and provides more nutrition availability for fungal hyphal development. As a result, the virulence of the engineered fungus against the insect pest was significantly improved compared to the wild-type strain. These findings indicate the significance of domain engineering in the development of more potent cuticle-degrading enzymes, providing a viable method to improve the biocontrol effectiveness of entomopathogenic fungi like *Metarhizium anisopliae* and *Beauveria bassiana* [19].

In the present study, we constructed chimeric cPr1A protease by the fusion of the *Bombyx mori* chitin-binding domain (BmChBD) to the C-terminus of the Pr1A protease from *Metarhizium anisopliae*. Our comparative binding and protease activity assay showed that the cPr1A revealed significantly improved substrate binding (28.9%, *p* < 0.002) and increased protease activity (35% increase, *p* < 0.03) compared to wild-type Pr1A. These results are consistent with those of Fan et al., who reported that the fusion of BmChBD to CDEP-1 enhanced cuticular chitin-binding, cuticle degradation, and pathogenicity via functional synergy between binding and cuticle degradation in entomopathogenic enzymes (19). The mutant fungi overexpressing the chimeric protease could accelerate host infection and decrease the time to kill insect pests, addressing one of the major drawbacks of *Metarhizium*-based biopesticides.

Nonetheless, the present research work is limited to in vitro assays, and further studies are required to evaluate the impact of the engineered protease on fungal virulence under in vivo and field conditions.

## Supporting information

Supplementary Data: Tables and DNA Sequences

## References

1. Vivekanandhan P, Swathy K, Bedini S, Shivakumar MS (2023) Bioprospecting of *Metarhizium anisopliae* derived crude extract: an eco-friendly insecticide against insect pests. Int J Trop Insect Sci 43:429–440. 10.1007/s42690-022-00935-y

2. De Sousa TK, Silva ATD, Soares FEDF (2025) Fungi-based bioproducts: a review in the context of One Health. Pathogens 14:463. 10.3390/pathogens14050463

3. Wend K, Zorrilla L, Freimoser FM, Gallet A (2024) Microbial pesticides – challenges and future perspectives for testing and safety assessment with respect to human health. Environ Health 23:49. 10.1186/s12940-024-01090-2

4. Peng ZY, Huang ST, Chen JT, et al. (2022) An update of a green pesticide: *Metarhizium anisopliae*. All Life 15:1141–1159. 10.1080/26895293.2022.2147224

5. Thavkar S, Thube S, Panchbhai P, et al. (2025) Evaluation of bio-efficacy of *Metarhizium anisopliae* against the pink bollworm *Pectinophora gossypiella* with insights into its colonization potential and insecticide compatibility. J Cotton Res 8:8. 10.1186/s42397-025-00213-5

6. Chowdhury MZH, Mostofa MG, Mim MF, et al. (2024) The fungal endophyte *Metarhizium anisopliae* (MetA1) coordinates salt tolerance mechanisms of rice to enhance growth and yield. Plant Physiol Biochem 207:108328. 10.1016/j.plaphy.2023.108328

7. Khachatourians GG, Uribe D (2004) Genomics of entomopathogenic fungi. In: Arora DK, Khachatourians GG (eds) *Fungal genomics*. Appl Mycol Biotechnol 4:353–378. 10.1016/S1874-5334(04)80018-2

8. Meyling NV, Eilenberg J (2007) Ecology of the entomopathogenic fungi *Beauveria bassiana* and *Metarhizium anisopliae* in temperate agroecosystems: potential for conservation biological control. Biol Control 43:145–155. 10.1016/j.biocontrol.2007.07.007

9. Zimmermann G (1993) The entomopathogenic fungus *Metarhizium anisopliae* and its potential as a biocontrol agent. J Pestic Sci 37:375–379. 10.1002/ps.2780370410

10. Zimmermann G (2007) Review on safety of the entomopathogenic fungus *Metarhizium anisopliae*. Biocontrol Sci Technol 17:879–920. 10.1080/09583150701593963

11. Keppanan R, Sivaperumal S, Ramos Aguila LC, et al. (2018) Isolation and characterization of *Metarhizium anisopliae* TK29 and its mycoinsecticide effects against subterranean termite *Coptotermes formosanus*. Microb Pathog 123:52–59. 10.1016/j.micpath.2018.06.040

12. Kaaya GP, Samish M, Hedimbi M, Gindin G, Glazer I (2011) Control of tick populations by spraying *Metarhizium anisopliae* conidia on cattle under field conditions. Exp Appl Acarol 55:273–281. 10.1007/s10493-011-9471-3

13. Butt TM, Greenfield BPJ, Greig C, et al. (2013) *Metarhizium anisopliae* pathogenesis of mosquito larvae: a verdict of accidental death. PLoS ONE 8:e81686. 10.1371/journal.pone.0081686

14. Leemon DM, Jonsson NN (2008) Laboratory studies on Australian isolates of *Metarhizium anisopliae* as a biopesticide for the cattle tick *Boophilus microplus*. J Invertebr Pathol 97:40–49. 10.1016/j.jip.2007.07.006

15. St Leger R, Joshi L, Bidochka MJ, Roberts DW (1996) Construction of an improved mycoinsecticide overexpressing a toxic protease. Proc Natl Acad Sci USA 93:6349–6354. 10.1073/pnas.93.13.6349

16. Bagga S, Hu G, Screen SE, St Leger RJ (2004) Reconstructing the diversification of subtilisins in the pathogenic fungus *Metarhizium anisopliae*. Gene 324:159–169. 10.1016/j.gene.2003.09.031

17. Leão MPC, Tiago PV, Andreote FD, Araújo WL, Oliveira NT (2015) Differential expression of the Pr1A gene in *Metarhizium anisopliae* and *Metarhizium acridum* across different culture conditions and during pathogenesis. Genet Mol Biol 38:86–92. 10.1590/S1415-475738138120140236

18. St Leger RJ, Joshi L, Roberts D (1998) Ambient pH is a major determinant in the expression of cuticle-degrading enzymes and hydrophobin by *Metarhizium anisopliae*. Appl Environ Microbiol 64:709–713. 10.1128/AEM.64.2.709-713.1998

19. Fan Y, Pei X, Guo S, et al. (2010) Increased virulence using engineered protease-chitin binding domain hybrid expressed in the entomopathogenic fungus *Beauveria bassiana*. Microb Pathog 49:376–380. 10.1016/j.micpath.2010.06.013

20. Joshi L, St Leger RJ, Roberts DW (1997) Isolation of a cDNA encoding a novel subtilisin-like protease (Pr1B) from the entomopathogenic fungus *Metarhizium anisopliae* using differential display-RT-PCR. Gene 197:1–8. 10.1016/S0378-1119(97)00132-7

21. Screen SE, Hu G, St Leger RJ (2001) Transformants of *Metarhizium anisopliae* sf. anisopliae overexpressing chitinase from *Metarhizium anisopliae* sf. acridum show early induction of native chitinase but are not altered in pathogenicity to *Manduca sexta*. J Invertebr Pathol 78:260–266. 10.1006/jipa.2001.5067

22. St Leger RJ, Joshi L, Bidochka MJ, Rizzo NW, Roberts DW (1996) Biochemical characterization and ultrastructural localization of two extracellular trypsins produced by *Metarhizium anisopliae* in infected insect cuticles. Appl Environ Microbiol 62:1257–1264. 10.1128/aem.62.4.1257-1264.1996

23. St Leger RJ, Joshi L, Bidochka MJ, Rizzo NW, Roberts DW (1996) Characterization and ultrastructural localization of chitinases from *Metarhizium anisopliae*, *M. flavoviride*, and *Beauveria bassiana* during fungal invasion of host (*Manduca sexta*) cuticle. Appl Environ Microbiol 62:907–912. 10.1128/aem.62.3.907-912.1996

24. Saciloto-de-Oliveira L, Innocente-Alves C, Sant’Ana JDF, et al. (2025) Proteomics in *Metarhizium* parasitism of arthropods. Fungal Biol Rev 51:100409. 10.1016/j.fbr.2024.100409

25. Schrank A, Vainstein MH (2010) *Metarhizium anisopliae* enzymes and toxins. Toxicon 56:1267–1274. 10.1016/j.toxicon.2010.03.008

26. Huang L, Wang Z, Davaasambuu U, et al. (2025) Effect of *Metarhizium anisopliae* IPPM202 extracellular proteinases on midgut of *Locusta migratoria manilensis*. Insects 16:1111. 10.3390/insects16111111

27. Svitil AL, Kirchman DL (1998) A chitin-binding domain in a marine bacterial chitinase and other microbial chitinases: implications for the ecology and evolution of 1,4-β-glycanases. Microbiology 144:1299–1308. 10.1099/00221287-144-5-1299

28. Ikegami T, Okada T, Hashimoto M, Seino S, Watanabe T, Shirakawa M (2000) Solution structure of the chitin-binding domain of *Bacillus circulans* WL-12 chitinase A1. J Biol Chem 275:13654–13661. 10.1074/jbc.275.18.13654

29. Arakane Y, Zhu Q, Matsumiya M, Muthukrishnan S, Kramer KJ (2003) Properties of catalytic, linker and chitin-binding domains of insect chitinase. Insect Biochem Mol Biol 33:631–648. 10.1016/S0965-1748(03)00049-3

30. Itoh Y, Kawase T, Nikaidou N, et al. (2002) Functional analysis of the chitin-binding domain of a family 19 chitinase from *Streptomyces griseus* HUT6037: substrate-binding affinity and cis-dominant increase of antifungal function. Biosci Biotechnol Biochem 66:1084–1092. 10.1271/bbb.66.1084

31. Iseli B, Boller T, Neuhaus JM (1993) The N-terminal cysteine-rich domain of tobacco class I chitinase is essential for chitin binding but not for catalytic or antifungal activity. Plant Physiol 103:221–226. 10.1104/pp.103.1.221

32. Taira T, Ohnuma T, Yamagami T, et al. (2002) Antifungal activity of rye (*Secale cereale*) seed chitinases: the different binding manner of class I and class II chitinases to the fungal cell walls. Biosci Biotechnol Biochem 66:970–977. 10.1271/bbb.66.970

33. Martínez-Zavala SA, Salcedo-Hernández R, Carballo-Uicab VM, Casados-Vázquez LE, Bideshi DK, Barboza-Corona JE (2025) Exposed tryptophan residues in the chitin-binding domain of ChiA74 chitinase are important for chitin-binding and antifungal activity. Int J Biol Macromol 302:140465. 10.1016/j.ijbiomac.2025.140465

34. Yu Y, Leng K, Cao R, et al. (2025) Functional roles of dual chitin-binding domains in *Chitiniphilus eburneus* chitinase properties. Int J Biol Macromol 329:147963. 10.1016/j.ijbiomac.2025.147963

35. Gu C, Chen J, Huang X, et al. (2025) The impact of chitinase binding domain truncation on the properties of CaChi18B from *Chitinilyticum aquatile* CSC-1. Mar Drugs 23:93. 10.3390/md23030093

36. Danielli A, Loukeris TG, Lagueux M, Müller HM, Richman A, Kafatos FC (2000) A modular chitin-binding protease associated with hemocytes and hemolymph in the mosquito *Anopheles gambiae*. Proc Natl Acad Sci USA 97:7136–7141. 10.1073/pnas.97.13.7136

37. Gorman MJ, Andreeva OV, Paskewitz SM (2000) Sp22D: a multidomain serine protease with a putative role in insect immunity. Gene 251:9–17. 10.1016/S0378-1119(00)00181-5

38. Miyamoto K, Nukui E, Itoh H, et al. (2002) Molecular analysis of the gene encoding a novel chitin-binding protease from *Alteromonas* sp. strain O-7 and its role in the chitinolytic system. J Bacteriol 184:1865–1872. 10.1128/JB.184.7.1865-1872.2002

39. Sidhu SS, Kalmar GB, Willis LG, Borgford TJ (1994) *Streptomyces griseus* protease C: a novel enzyme of the chymotrypsin superfamily. J Biol Chem 269:20167–20171. 10.1016/S0021-9258(17)32141-5

40. Image 1 - Pictorial illustration of construction and cloning of chimeric cPr1A gene into pET-28a vector.

