## Supplementary Data: Tables and DNA Sequences for "Bioengineering of a Chimeric *Metarhizium anisopliae* cPr1A Protease with Enhanced Binding and Enzyme Activity Through C-terminal Fusion of *Bombyx mori* Chitin-Binding Domain"

**Table S1: Composition of the laboratory-prepared Gibson Assembly Master Mix**

| **Component** | **Volume (µl)** |
| --- | --- |
| 5× Isothermal Mix | 30.8 |
| T5 Exonuclease (10 U/µl) | 0.4 |
| Phusion High-Fidelity DNA Polymerase (2 U/µl) | 1.2 |
| Taq DNA Ligase (40 U/µl) | 1.6 |
| Nuclease-free water | 86.0 |
| Total volume | 120.0 |

**Table S2: PCR cycles conditions for amplification of Pr1A from *Metarhizium anisopliae***

| **Steps** |  | **Time** |
| --- | --- | --- |
|  |  | 2 min |
| I. Denaturation at 98°C |  |  |
| II. 30 cycles of | i) Denaturation at 98°C | 10 s |
|  | ii) Annealing at 68°C | 30 s |
|  | iii) Extension at 72°C | 30 s |
| III. Final extension at 72°C |  | 10 min |

**Table S3: Primer sequences used for amplification of Pr1A gene and Pr1A-Bm gene fragment from *Metarhizium anisopliae* and Bm-Pr1A gene fragment from *Bombyx mori***

| **Gene construct** | **Primer Sequence** |
| --- | --- |
| ***Pr1A_FP_NcoI_pET28a_Overhang*** | CTTTAAGAAGGAGATATACCATG GGCATCACTGAGCAGAGCGG |
| ***Pr1A_RP_His Tag_XhoI_pET28a_Overhang*** | GGTGGTGGTGGTGCTCGAGTTAATGATGATGATGATGATGGGCACCGTTG TAGGCAA |
| ***Pr1A-Bm_FP_NcoI_pET28a_Overhang*** | CTTTAAGAAGGAGATATACCATG GGCATCACTGAGCAGAGCGG |
| ***Pr1A-Bm _RP_BmChBD_Overhang*** | CCTCAGAGTTACACACATC GGCACCGTTGTAGGCAAGGTAG |
| ***Bm-Pr1A_FP_Pr1A_Overhang*** | TTGCCTACAACGGTGCC GATGTGTGTAACTCTGAGG |
| ***Bm-Pr1A_RP_His Tag_XhoI_pET28a_Overhang*** | GGTGGTGGTGGTGCTCGAGTTAATGATGATGATGATGATGAGGCCAATCGCAAACGTTAAG |

**Table S4: PCR cycles conditions for amplification of Pr1A-Bm from *Metarhizium anisopliae***

| **Steps** |  | **Time** |
| --- | --- | --- |
|  |  | 2 min |
| I. Denaturation at 98°C |  |  |
| II. 30 cycles of | i) Denaturation at 98°C | 10 s |
|  | ii) Annealing at 68°C | 30 s |
|  | iii) Extension at 72°C | 45 s |
| III. Final extension at 72°C |  | 10 min |

**Table S5: PCR cycles conditions for amplification of Bm-Pr1A from *Bombyx mori***

| **Steps** |  | **Time** |
| --- | --- | --- |
|  |  | 2 min |
| I. Denaturation at 98°C |  |  |
| II. 30 cycles of | i) Denaturation at 98°C | 10 s |
|  | ii) Annealing at 70°C | 30 s |
|  | iii) Extension at 72°C | 15 s |
| III. Final extension at 72°C |  | 10 min |

**Table S6: Specific activity comparison of purified wild-type and chimeric Pr1A protease**

| **Enzyme** | **Specific Activity (U/mg)** | **Relative Activity (%)** |
| --- | --- | --- |
| Wild-type Pr1A | 0.254 ± 0.06 | 100 |
| Chimeric cPr1A | 0.343 ± 0.08 | 135 |

**Table S7: Secondary structure analysis of Pr1A and cPr1A proteins**

| **Protein** | **α helix** | **β strands** | **Random coils** |
| --- | --- | --- | --- |
| Pr1A | 25.4 % | 17.99 % | 56.6 % |
| cPr1A | 21.7 % | 18.34 % | 59.9 % |

***Metarhizium anisopliae* Pr1A gene DNA sequence GenBank: M73795.1**

GGCATCACTGAGCAGAGCGGTGCTCCCTGGGGTCTTGGGCGCATCTCTCACCGCAGTAAGGGAAGCACCACCTATCGCTACGATGATAGTGCTGGTCAGGGTACTTGCGTATATATCATTGACACTGGTATTGAGGCCTCCCACCCCGAGTTTGAGGGTCGCGCCACTTTTCTTAAGAGCTTCATCAGCGGTCAAAACACTGATGGCCACGGCCATGGGACTCACTGCGCTGGTACCATTGGTAGCAAGACCTACGGTGTTGCCAAAAAGGCTAAGCTCTATGGTGTCAAGGTTCTTGACAACCAGGGCAGTGGTTCCTACTCCGGTATCATCAGTGGCATGGACTACGTTGCACAGGACTCCAAGACCCGCGGCTGCCCCAACGGCGCCATTGCTTCCATGAGCCTGGGAGGTGGCTACTCGGCGTCCGTCAACCAAGGTGCTGCTGCTTTGGTCAATTCTGGTGTCTTCCTTGCCGTCGCCGCTGGCAACGATAACCGGGATGCCCAGAACACCTCTCCCGCTTCCGAGCCTTCTGCCTGCACTGTTGGTGCCTCTGCGGAAAATGACAGCCGATCTTCCTTCTCCAACTACGGCAGAGTTGTCGATATTTTCGCTCCTGGTAGCAATGTTCTTTCCACCTGGATTG**T**TGGCCGCACAAAC**T**CCATCTCTGGTACCTCCATGGCTACTCCCCATATTGCCGGTCTGGCTGCCTACCTCAGTGCGCTCCAAGGCAAGACTACCCCTGCCGCTCTTTGCAAGAAGATCCAGGACACTGCTACCAAGAACGTGCTCACCGGTGTTCCCTCTGGCACTGTCAACTACCTTGCCTACAACGGTGCCTAA

**Supplementary Figure S1.** Pr1A DNA sequence retrieved from NCBI (accession no. M73795.1).

**The deduced sequence of Pr1A (*Metarhizium anisopliae* 892)**

GGCATCACTGAGCAGAGCGGTGCTCCCTGGGGTCTTGGGCGCATCTCTCACCGCAGTAAGGGAAGCACCACCTATCGCTACGATGATAGTGCTGGTCAGGGTACTTGCGTATATATCATTGACACTGGTATTGAGGCCTCCCACCCCGAGTTTGAGGGTCGCGCCACTTTTCTTAAGAGCTTCATCAGCGGTCAAAACACTGATGGCCACGGCCATGGGACTCACTGCGCTGGTACCATTGGTAGCAAGACCTACGGTGTTGCCAAAAAGGCTAAGCTCTATGGTGTCAAGGTTCTTGACAACCAGGGCAGTGGTTCCTACTCCGGTATCATCAGTGGCATGGACTACGTTGCACAGGACTCCAAGACCCGCGGCTGCCCCAACGGCGCCATTGCTTCCATGAGCCTGGGAGGTGGCTACTCGGCGTCCGTCAACCAAGGTGCTGCTGCTTTGGTCAATTCTGGTGTCTTCCTTGCCGTCGCCGCTGGCAACGATAACCGGGATGCCCAGAACACCTCTCCCGCTTCCGAGCCTTCTGCCTGCACTGTTGGTGCCTCTGCGGAAAATGACAGCCGATCTTCCTTCTCCAACTACGGCAGAGTTGTCGATATTTTCGCTCCTGGTAGCAATGTTCTTTCCACCTGGATTG**G**TGGCCGCACAAAC**A**CCATCTCTGGTACCTCCATGGCTACTCCCCATATTGCCGGTCTGGCTGCCTACCTCAGTGCGCTCCAAGGCAAGACTACCCCTGCCGCTCTTTGCAAGAAGATCCAGGACACTGCTACCAAGAACGTGCTCACCGGTGTTCCCTCTGGCACTGTCAACTACCTTGCCTACAACGGTGCCTAA

**Supplementary Figure S2.** Deduced sequence of the mature Pr1A domain from *Metarhizium anisopliae* strain 892 used in this study.

**Multiple sequence alignment of Pr1A DNA sequences**

```
1      GGCATCACTGAGCAGAGCGGTGCTCCCTGGGGTCTTGGGCGCATCTCTCACCGCAGTAAG	60
2      GGCATCACTGAGCAGAGCGGTGCTCCCTGGGGTCTTGGGCGCATCTCTCACCGCAGTAAG	60
       ************************************************************

1      GGAAGCACCACCTATCGCTACGATGATAGTGCTGGTCAGGGTACTTGCGTATATATCATT	120
2      GGAAGCACCACCTATCGCTACGATGATAGTGCTGGTCAGGGTACTTGCGTATATATCATT	120
       ************************************************************

1      GACACTGGTATTGAGGCCTCCCACCCCGAGTTTGAGGGTCGCGCCACTTTTCTTAAGAGC	180
2      GACACTGGTATTGAGGCCTCCCACCCCGAGTTTGAGGGTCGCGCCACTTTTCTTAAGAGC	180
       ************************************************************

1      TTCATCAGCGGTCAAAACACTGATGGCCACGGCCATGGGACTCACTGCGCTGGTACCATT	240
2      TTCATCAGCGGTCAAAACACTGATGGCCACGGCCATGGGACTCACTGCGCTGGTACCATT	240
       ************************************************************

1      GGTAGCAAGACCTACGGTGTTGCCAAAAAGGCTAAGCTCTATGGTGTCAAGGTTCTTGAC	300
2      GGTAGCAAGACCTACGGTGTTGCCAAAAAGGCTAAGCTCTATGGTGTCAAGGTTCTTGAC	300
       ************************************************************

1      AACCAGGGCAGTGGTTCCTACTCCGGTATCATCAGTGGCATGGACTACGTTGCACAGGAC	360
2      AACCAGGGCAGTGGTTCCTACTCCGGTATCATCAGTGGCATGGACTACGTTGCACAGGAC	360
       ************************************************************

1      TCCAAGACCCGCGGCTGCCCCAACGGCGCCATTGCTTCCATGAGCCTGGGAGGTGGCTAC	420
2      TCCAAGACCCGCGGCTGCCCCAACGGCGCCATTGCTTCCATGAGCCTGGGAGGTGGCTAC	420
       ************************************************************

1      TCGGCGTCCGTCAACCAAGGTGCTGCTGCTTTGGTCAATTCTGGTGTCTTCCTTGCCGTC	480
2      TCGGCGTCCGTCAACCAAGGTGCTGCTGCTTTGGTCAATTCTGGTGTCTTCCTTGCCGTC	480
       ************************************************************

1      GCCGCTGGCAACGATAACCGGGATGCCCAGAACACCTCTCCCGCTTCCGAGCCTTCTGCC	540
2      GCCGCTGGCAACGATAACCGGGATGCCCAGAACACCTCTCCCGCTTCCGAGCCTTCTGCC	540
       ************************************************************

1      TGCACTGTTGGTGCCTCTGCGGAAAATGACAGCCGATCTTCCTTCTCCAACTACGGCAGA	600
2      TGCACTGTTGGTGCCTCTGCGGAAAATGACAGCCGATCTTCCTTCTCCAACTACGGCAGA	600
       ************************************************************

1      GTTGTCGATATTTTCGCTCCTGGTAGCAATGTTCTTTCCACCTGGATTGTTGGCCGCACA	660
2      GTTGTCGATATTTTCGCTCCTGGTAGCAATGTTCTTTCCACCTGGATTGGTGGCCGCACA	660
       ************************************************* **********

1      AACTCCATCTCTGGTACCTCCATGGCTACTCCCCATATTGCCGGTCTGGCTGCCTACCTC	720
2      AACACCATCTCTGGTACCTCCATGGCTACTCCCCATATTGCCGGTCTGGCTGCCTACCTC	720
       *** ********************************************************

1      AGTGCGCTCCAAGGCAAGACTACCCCTGCCGCTCTTTGCAAGAAGATCCAGGACACTGCT	780
2      AGTGCGCTCCAAGGCAAGACTACCCCTGCCGCTCTTTGCAAGAAGATCCAGGACACTGCT	780
       ************************************************************

1      ACCAAGAACGTGCTCACCGGTGTTCCCTCTGGCACTGTCAACTACCTTGCCTACAACGGT	840
2      ACCAAGAACGTGCTCACCGGTGTTCCCTCTGGCACTGTCAACTACCTTGCCTACAACGGT	840
       ************************************************************

1      GCCTAA		846
2      GCCTAA		846
       ******
```

**Supplementary Figure S3.** Multiple sequence alignment of the Pr1A DNA sequence. Sequence 1: Pr1A DNA sequence retrieved from NCBI (accession no. M73795.1). Sequence 2: Deduced sequence of the mature Pr1A domain from *Metarhizium anisopliae* strain 892 used in this study (signal peptide and propeptide excluded). Nucleotide differences in the mature domain are highlighted.

**Multiple sequence alignment of Pr1A amino acid sequences**

```
1      GITEQSGAPWGLGRISHRSKGSTTYRYDDSAGQGTCVYIIDTGIEASHPEFEGRATFLKS	60
2      GITEQSGAPWGLGRISHRSKGSTTYRYDDSAGQGTCVYIIDTGIEASHPEFEGRATFLKS	60
       ************************************************************

1      FISGQNTDGHGHGTHCAGTIGSKTYGVAKKAKLYGVKVLDNQGSGSYSGIISGMDYVAQD	120
2      FISGQNTDGHGHGTHCAGTIGSKTYGVAKKAKLYGVKVLDNQGSGSYSGIISGMDYVAQD	120
       ************************************************************

1      SKTRGCPNGAIASMSLGGGYSASVNQGAAALVNSGVFLAVAAGNDNRDAQNTSPASEPSA	180
2      SKTRGCPNGAIASMSLGGGYSASVNQGAAALVNSGVFLAVAAGNDNRDAQNTSPASEPSA	180
       ************************************************************

1      CTVGASAENDSRSSFSNYGRVVDIFAPGSNVLSTWIVGRTNSISGTSMATPHIAGLAAYL	240
2      CTVGASAENDSRSSFSNYGRVVDIFAPGSNVLSTWIGGRTNTISGTSMATPHIAGLAAYL	240
       ************************************ ****:******************

1      SALQGKTTPAALCKKIQDTATKNVLTGVPSGTVNYLAYNGA		281
2      SALQGKTTPAALCKKIQDTATKNVLTGVPSGTVNYLAYNGA		281
       *****************************************
```

**Supplementary Figure S4.** Multiple sequence alignment of Pr1A amino acid sequences. Sequence 1: Pr1A from NCBI reference (accession no. M73795.1). Sequence 2: Deduced amino acid sequence of the mature Pr1A domain from *Metarhizium anisopliae* strain 892 used in this study (signal peptide and propeptide excluded, residue numbering starting from position 1 at the N-terminal catalytic residue). Amino acid differences in the mature domain are highlighted.
